# Robust designation of meiotic crossover sites by CDK-2 through phosphorylation of the MutSγ complex

**DOI:** 10.1101/2021.08.31.458431

**Authors:** Jocelyn Haversat, Alexander Woglar, Kayla Klatt, Chantal C. Akerib, Victoria Roberts, Catcher C. Salazar, Shin-Yu Chen, Swathi Arur, Anne M. Villeneuve, Yumi Kim

## Abstract

Crossover formation is essential for proper segregation of homologous chromosomes during meiosis. Here we show that *C. elegans* Cyclin-dependent kinase 2 (CDK-2) forms a complex with cyclin-like protein COSA-1 and supports crossover formation by promoting conversion of meiotic double-strand breaks (DSBs) into crossover-specific recombination intermediates. Further, we identify MutSγ component MSH-5 as a CDK-2 phosphorylation target. MSH-5 has a disordered C-terminal tail that contains 13 potential CDK phosphosites and is required to concentrate crossover-promoting proteins at recombination sites. Phosphorylation of the MSH-5 tail appears dispensable in a wild-type background, but when MutSγ activity is partially compromised, crossover formation and retention of CDK-2/COSA-1 at recombination sites are exquisitely sensitive to phosphosite loss. Our data support a model in which robustness of crossover designation reflects a positive feedback mechanism involving CDK-2-mediated phosphorylation and scaffold-like properties of the MSH-5 C-terminal tail, features that combine to promote full recruitment and activity of crossover-promoting complexes.

## INTRODUCTION

Sexually reproducing organisms rely on proper chromosome segregation during meiosis to produce gametes with a complete genome. During meiotic prophase I, chromosomes pair and undergo crossover recombination with their homologous partners. This process, together with sister chromatid cohesion, leads to the formation of physical linkages between the homologs and enables their separation during meiosis I. Defects in crossover formation are disastrous, leading to miscarriages and congenital disorders, such as Down Syndrome (MacLennan et al., 2015).

Meiotic recombination initiates with the generation of programmed DNA double-strand breaks (DSBs) by the topoisomerase-like enzyme Spo11 (Keeney et al., 1997). DSBs are resected to yield two 3’ end single-stranded DNA (ssDNA) overhangs, which are rapidly coated by RecA recombinases, Dmc1 and Rad51. This nucleoprotein filament then seeks out homology and invades a homologous template, forming a metastable single-end invasion intermediate (D-loop) (Hunter and Kleckner, 2001). If the extended D-loop is captured by ssDNA on the other side of the DSB in a process known as second-end capture, a double Holliday junction (dHJ) forms (Schwacha and Kleckner, 1995). During meiosis, dHJs are specifically resolved as crossovers by the endonuclease MutLγ (MLH1-MLH3) (Allers and Lichten, 2001; Marsolier-Kergoat et al., 2018; Zakharyevich et al., 2012). Although a multitude of DSBs are generated during meiotic prophase, strikingly few are ultimately selected to become crossovers. Early recombination intermediates pare down in pachytene until each homolog pair receives at least one crossover, while the majority of DSBs are repaired as non-crossovers (Gray and Cohen, 2016). However, how meiotic DSBs are chosen to become crossovers remains poorly understood.

Throughout eukaryotes, crossover recombination is primarily controlled by a group of proteins collectively termed “ZMM” (Pyatnitskaya et al., 2019). Notably, homologs of the yeast Ring domain protein Zip3 (ZHP-1, -2, -3, and -4 in *C. elegans* (Bhalla et al., 2008; Jantsch et al., 2004; Nguyen et al., 2018; Zhang et al., 2018), *Drosophila* Vilya and Narya/Nenya (Lake et al., 2019; Lake et al., 2015), Hei10 in *Arabidopsis* (Chelysheva et al., 2012), and Hei10 and RNF212 in mammals (Reynolds et al., 2013; Ward et al., 2007)) initially localize as abundant foci or long stretches along the synaptonemal complex (SC) but eventually concentrate at crossover sites in late pachytene (Agarwal and Roeder, 2000). These SUMO or ubiquitin ligases appear to promote crossover designation by stabilizing the ZMM proteins at crossover sites while removing them from other recombination intermediates (Nguyen et al., 2018; Qiao et al., 2014; Rao et al., 2017; Reynolds et al., 2013; Ward et al., 2007; Zhang et al., 2018). Although many meiotic proteins are shown to be SUMO-modified (Bhagwat et al., 2021), key targets of the Zip3 family proteins remain largely unknown.

The meiosis-specific MutS homologs, MSH4 and MSH5, form a heterodimeric MutSγ complex and play essential roles in crossover formation in diverse eukaryotes (de Vries et al., 1999; Higgins et al., 2004; Higgins et al., 2008; Hollingsworth et al., 1995; Kelly et al., 2000; Milano et al., 2019; Ross-Macdonald and Roeder, 1994; Zalevsky et al., 1999). MutSγ localizes to recombination intermediates as numerous foci but ultimately accumulates at sites that are destined to become crossovers (Kneitz et al., 2000; Woglar and Villeneuve, 2018). Biochemical analyses using recombinant MSH4 and MSH5 have shown that MutSγ recognizes single-end invasion intermediates and Holliday junctions (HJs) *in vitro* (Lahiri et al., 2018; Snowden et al., 2004). HJs activate the ATP hydrolysis of MutSγ and promote the exchange of bound ADP for ATP, inducing the formation of a sliding clamp that dissociates from HJs (Snowden et al., 2004; Snowden et al., 2008). By iterative loading and embracing DNA duplexes within a dHJ, MutSγ is thought to stabilize crossover-specific recombination intermediates (Snowden et al., 2004; Woglar and Villeneuve, 2018). In addition, MutSγ recruits and activates the resolvase activity of MutLγ, enabling biased processing of dHJs into crossovers during meiosis (Cannavo et al., 2020; Kulkarni et al., 2020).

A genetic screen in *C. elegans* has identified a cyclin-like protein COSA-1 as a component essential for processing meiotic DSBs into crossovers (Yokoo et al., 2012). The mammalian ortholog CNTD1 was subsequently identified (Holloway et al., 2014), and both COSA-1 and CNTD1 have been shown to localize to crossover sites (Bondarieva et al., 2020; Gray et al., 2020; Yokoo et al., 2012). In the absence of COSA-1/CNTD1, MutSγ components persist as numerous foci in pachytene, and crossover formation is eliminated or severely compromised (Holloway et al., 2014; Woglar and Villeneuve, 2018), demonstrating a crucial role of COSA-1/CNTD1 in converting early recombination intermediates into crossovers. Since both COSA-1 and CNTD1 are members of the cyclin family, it is plausible that they form a complex with a cyclin-dependent kinase (CDK) and regulate the recombination process through phosphorylation.

Several lines of evidence suggest that CDK2 might be the kinase partner for CNTD1. CDK2 interacts with CNTD1 in yeast-two hybrid assays (Bondarieva et al., 2020; Gray et al., 2020) and localizes to interstitial chromosome sites (Ashley et al., 2001; Liu et al., 2014) in a CNTD1-dependent manner (Holloway et al., 2014). Reduced CDK2 activity leads to a failure in crossover formation, while a hyperactive form of CDK2 causes an increased number of MLH1 foci (Palmer et al., 2020). However, due to its requirement at telomeres in tethering chromosomes to the nuclear envelope, deletion of CDK2 leads to severe defects in SC assembly between paired homologs (synapsis) and pachytene arrest (Berthet et al., 2003; Ortega et al., 2003; Viera et al., 2015; Viera et al., 2009). Thus, it has been difficult to determine the role of CDK2 in crossover recombination. Moreover, key meiotic targets of CDK2 have not yet been identified.

We reasoned that CDK-2, the *C. elegans* homolog of CDK2, might also localize and function at crossover sites. However, global knockdown of *C. elegans* CDK-2 by RNAi leads to cell cycle arrest of mitotically proliferating germ cells (Jeong et al., 2011), thereby precluding the analysis of its requirement during meiotic prophase. To overcome this limitation and establish the meiotic function of CDK-2, we employ the auxin-inducible degradation system to deplete CDK-2 from the adult germline, demonstrating that CDK- 2 partners with COSA-1 to promote crossover formation during *C. elegans* meiosis. Moreover, we identify MSH-5 as a key substrate for CDK-2 and provide evidence that CDK-2/COSA-1 promotes crossover designation through phosphorylation and activation of the MutSγ complex.

## RESULTS

### CDK-2 colocalizes with COSA-1 both at early recombination intermediates and at crossover sites in pachytene

To establish the localization and function of CDK-2 during meiosis, we used CRISPR genome editing to generate a worm strain expressing CDK-2 fused to an auxin-inducible degron (AID) and three tandem Flag epitopes (3xFlag). Self-progeny of this worm strain are fully viable (100% egg viability), indicating that the AID tag does not interfere with essential CDK-2 functions. Immunofluorescence in whole-mount adult hermaphrodite germlines (XX) revealed that CDK-2 localizes to six distinct foci per nucleus in late pachytene, which correspond to crossover-designated recombination sites as marked by COSA-1 (**Figures 1A and S1A**). In male germlines (XO), CDK-2 appears as five foci in pachytene nuclei **(Figure S1B)**, consistent with its localization to crossover sites on the five autosomes.

**Figure 1.**
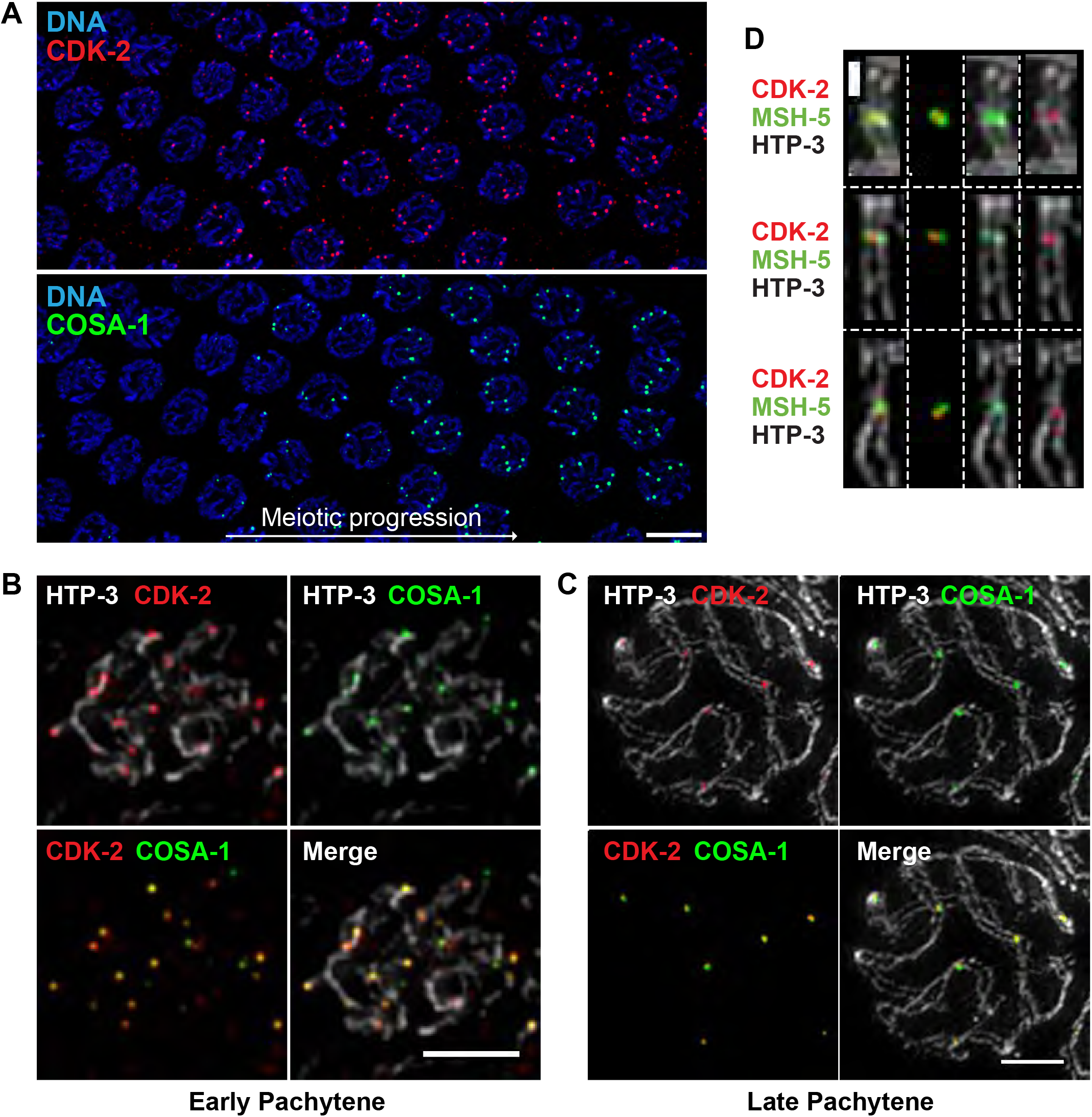
CDK-2 colocalizes with COSA-1 both at early recombination intermediates and at late pachytene crossovers. (A) Immunofluorescence images of a whole mount gonad from a worm strain that expresses CDK-2::AID::3Flag and GFP::COSA-1; the region of the gonad depicted shows CDK-2 and COSA-1 colocalizing in 6 bright foci in nuclei upon transition to late pachytene. Scale bar, 5 μm. (B-C) Full projections of Structured Illumination Microscopy (SIM) images of spread gonads showing the staining for HTP-3 (white), CDK-2 (red) and COSA- 1 (green) from early pachytene (B) and late pachytene nuclei (C). Scale bars, 2 μm. (D) Representative SIM images of individual crossover-designated sites showing CDK-2 singlet foci (red) localizing together with MSH-5 doublets (green). Scale bar, 500 nm.

We employed nuclear spreading and 3D structured illumination microscopy (SIM) to examine the localization of CDK-2 in relation to other crossover factors during meiotic progression. This cytological approach shows COSA-1 localizing to numerous early recombination intermediates as faint foci prior to transition to late pachytene (Woglar and Villeneuve, 2018). We likewise detected CDK-2 as numerous foci in early pachytene nuclei, colocalizing with COSA-1 **(Figure 1B)**. Upon transition to late pachytene, CDK-2 and COSA-1 are lost from most recombination sites and enriched together at six crossover-designated sites **(Figure 1C)**. Recent evidence has further shown that a distinct substructure emerges at the crossover site, in which a MSH-5 doublet appears orthogonal to chromosome axes, flanking a central COSA-1 focus (Woglar and Villeneuve, 2018). CDK-2 was similarly detected as a single focus positioned between two MSH-5 foci at the crossover site (**Figure 1D)**. Thus, CDK-2 colocalizes with COSA-1 at early recombination intermediates as well as crossover-designated sites in both hermaphrodite and male germlines.

### CDK-2 is required for crossover formation

To assess CDK-2 function during meiosis, we generated a strain in which CDK- 2::AID::3xFlag is expressed in conjunction with germline expressed plant F-box protein TIR1 (Zhang et al., 2015) (**Figure 2A**). Within 6 hours of 1 mM auxin treatment, CDK-2 was no longer detected in pachytene nuclei by immunofluorescence, demonstrating its rapid and efficient degradation (**Figure 2B**). The signal for COSA-1 was also completely lost in CDK-2-depleted germlines, indicating that COSA-1 localizes to recombination sites in a CDK-2-dependent manner.

**Figure 2.**
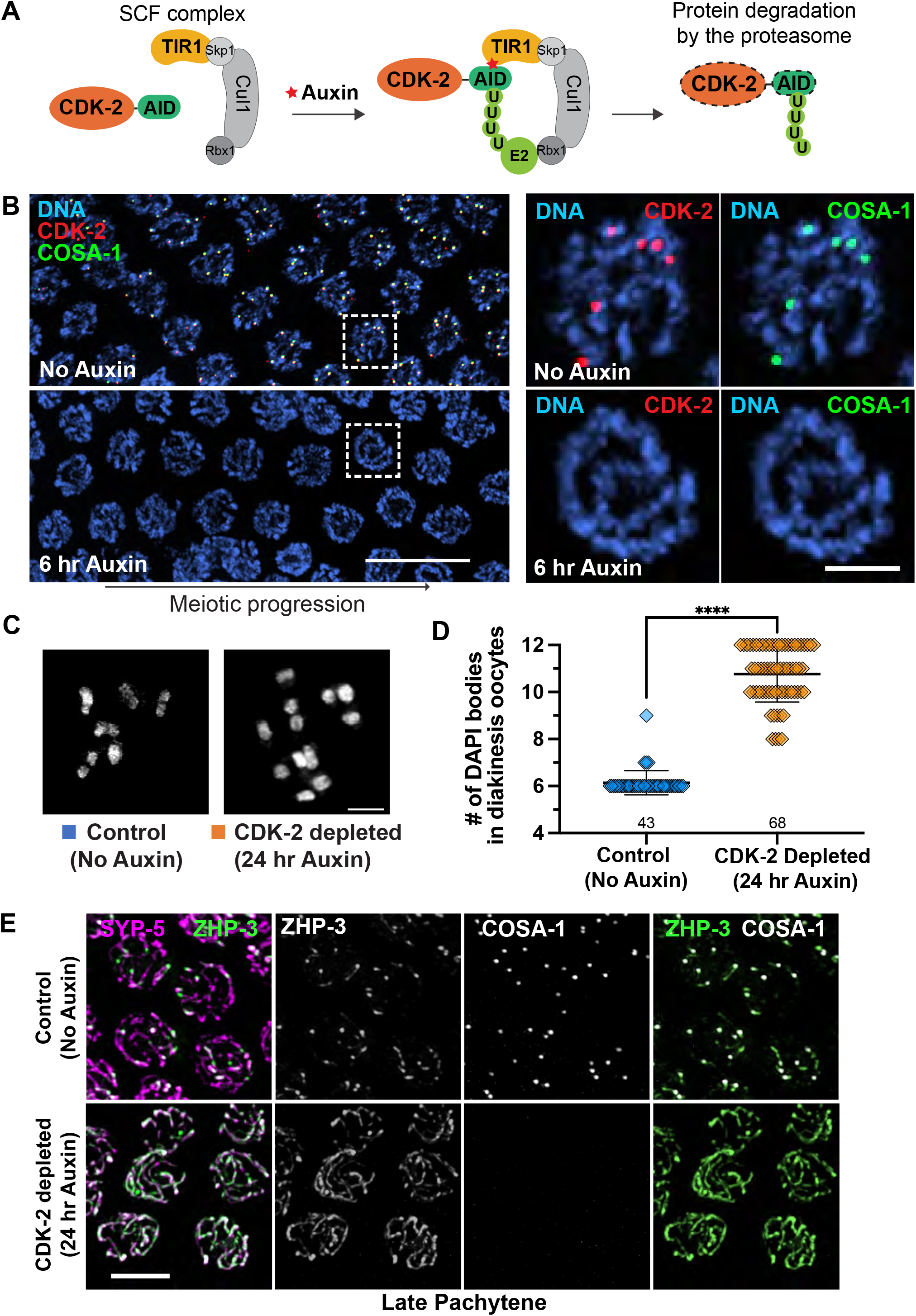
CDK-2 is required for crossover formation. (A) Schematic showing the auxin-inducible degradation system for CDK-2. TIR1::mRuby expressed from the *sun-1* promoter (*psun-1*) enables auxin-regulated control of CDK-2 degradation specifically in the germline. (B) Left: Young adult hermaphrodites expressing CDK-2::AID::3xFlag, GFP::COSA-1, and *Psun-1*::TIR1::mRuby were treated with 1 mM auxin for indicated times. Immunofluorescence images of late pachytene nuclei are shown. Scale bar, 10 μm. Right: Pachytene nuclei from the boxed regions at the indicated time points after 1 mM auxin treatment. Scale bar, 2 μm. (C) DAPI-stained oocyte nuclei in diakinesis from control (no auxin) and CDK-2-depleted animals (24 hr auxin), fixed and stained at 24 hr post L4. Scale bar, 3 μm. (D) Graph showing the number of DAPI bodies at diakinesis. ****, p<0.0001 by Mann-Whitney test. (E) Young adult hermaphrodites expressing CDK-2::AID::3xFlag, GFP::COSA-1, ZHP-3::V5, and TIR1::mRuby were treated with or without 1 mM auxin for 24 hr post L4 and dissected for immunofluorescence. Late pachytene nuclei stained for ZHP-3::V5 (green), SYP-5 (magenta), and GFP::COSA-1 (white) are shown. Scale bar, 4 μm.

Depletion of CDK-2 resulted in loss of crossover-based connections (chiasmata) that maintain associations between homologs in oocytes at diakinesis, the last stage of meiotic prophase. Six DAPI-stained bodies corresponding to six pairs of homologs connected by chiasmata (bivalents) are observed in wild-type diakinesis oocytes. However, following 24-hour auxin treatment, CDK-2-depleted oocytes displayed 10-12 DAPI-stained bodies, reflecting failure to form crossovers **(Figures 2C-D)**. Further, the RING domain-containing protein ZHP-3, which normally becomes restricted to six crossover sites in late pachytene nuclei (Bhalla et al., 2008), failed to become restricted to foci in CDK-2-depleted gonads and instead persisted along the SC **(Figure 2E)**, reflecting a requirement for CDK-2 in crossover formation.

Consistent with previous studies implicating *C. elegans* CDK-2 in the mitosis-to-meiosis decision and in promoting the proliferative fate of germline stem cells (Fox et al., 2011; Jeong et al., 2011), depletion of CDK-2 by 24-hr auxin treatment dramatically reduced the number of germ cells in the premeiotic region of the gonad **(Figures S2A-B)**. However, these CDK-2-depleted gonads exhibited normal pairing of HIM-8, a protein that binds a special region on X chromosomes known as the pairing center (Phillips et al., 2005), and robust synapsis **(Figures S2C-D)**. The sole RecA recombinase in *C. elegans*, RAD-51 (Alpi et al., 2003; Colaiacovo et al., 2003), was also detected as numerous foci in both control and CDK-2-depleted gonads **(Figure S2E)**. Thus, meiotic DSBs are induced in CDK-2-depleted germlines but cannot be processed into crossovers in the absence of CDK-2.

### CDK-2 is required for formation or stabilization of crossover-specific recombination intermediates

Next we used a partial nuclear spreading protocol that maintains the temporal and spatial organization of the gonad (Pattabiraman et al., 2017; Woglar and Villeneuve, 2018) to visualize the effects of CDK-2 depletion on the progression and architecture of meiotic recombination sites. To this end, we tagged MSH-5 at its C-terminus with the 14 amino acid long V5 epitope in a strain expressing CDK-2::AID::3xFlag and TIR1::mRuby and visualized MSH-5::V5 together with DSB-2, a marker of early meiotic prophase (Rosu et al., 2013) and an axis component, HTP-3. As previously demonstrated (Woglar and Villeneuve, 2018; Yokoo et al., 2012), MSH-5 localized as numerous foci corresponding to nascent recombination sites in early pachytene nuclei and ultimately pared down to six robust crossover-site foci in late pachytene in controls **(Figures 3A-B)**. In CDK-2- depleted germlines, however, MSH-5 persisted longer as multiple foci, and the early pachytene region positive for DSB-2 staining was extended **(Figures 3A-B)**, reflecting delayed meiotic progression due to failure to form crossover intermediates (Rosu et al., 2013; Stamper et al., 2013). Further, although CDK-2-depleted germ cells did eventually lose the DSB-2 signal and transition into the late pachytene stage, multiple faint MSH-5 foci persisted along chromosome axes **(Figure 3B)**, implying a failure of crossover designation in the absence of CDK-2. We note that abnormally large puncta of MSH-5 (white arrowheads) were detected in late pachytene nuclei of CDK-2-depleted germlines, which likely represent pathological aggregates of MSH-5 formed in the absence of CDK- 2 (See Discussion).

**Figure 3.**
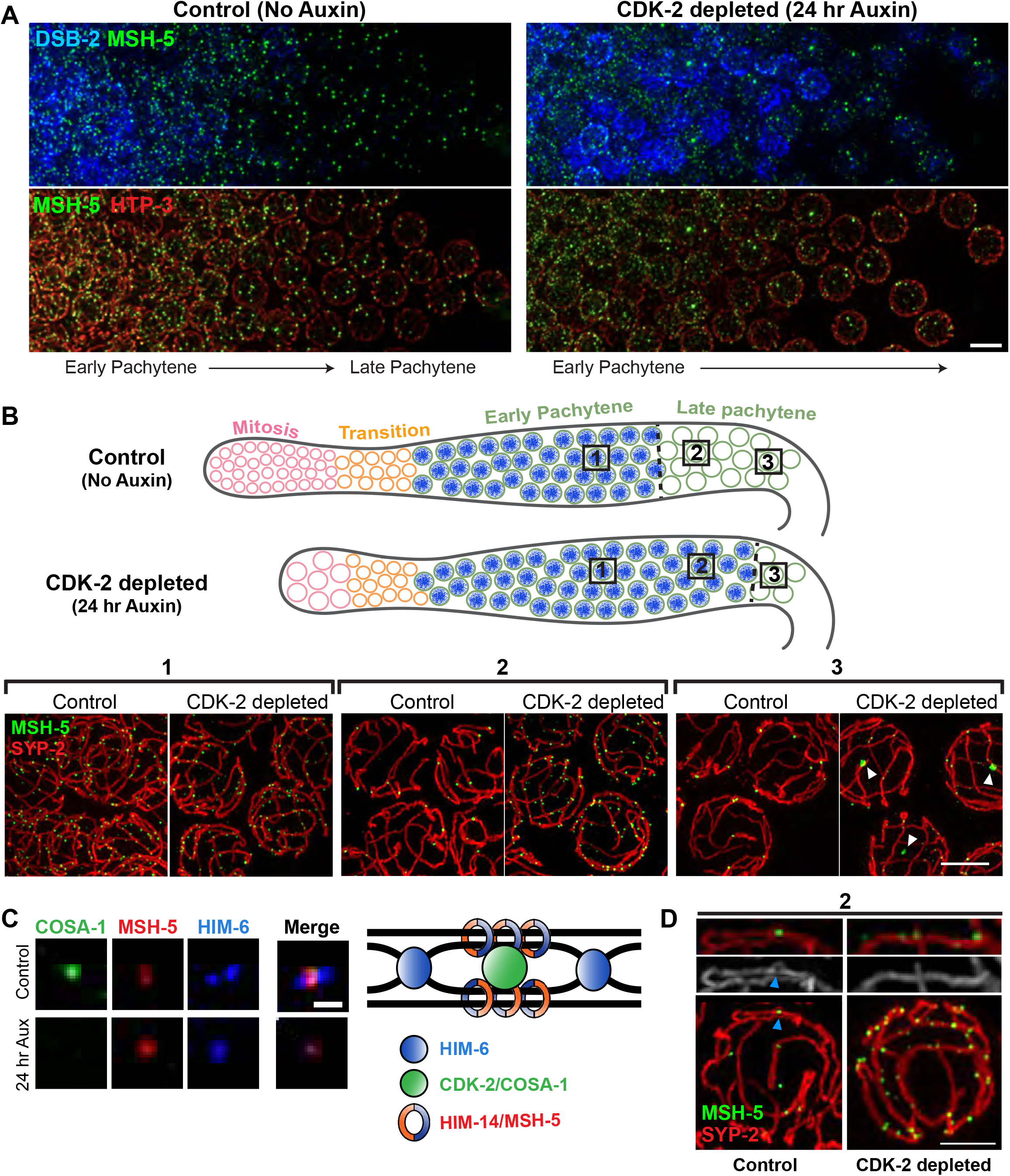
CDK-2 is required for stabilizing crossover-specific recombination intermediates. (A) Animals expressing CDK-2::AID::3xFlag and TIR1::mRuby were treated with or without 1 mM auxin for 24 hr post L4. Dissected gonads were spread and stained for DSB-2 (blue), MSH-5 (green), and HTP-3 (red). Scale bar, 5 μm. (B) Top: Diagram illustrating the effect of CDK-2 depletion on meiotic progression. The DSB-2 positive nuclei (shown in blue) represent nuclei in early pachytene. Bottom: Representative SIM images of nuclei from the indicated regions (1, 2 or 3) of spread gonads from control and CDK-2-depleted worms as indicated in the diagram above. White arrowheads in (3) indicate large MSH-5 aggregates. Scale bar, 3 μm. (C) Representative fluorescent images of recombination sites in late pachytene from control vs. CDK-2 depleted germline. Staining for MSH-5, HIM-6, and COSA-1 are shown. Scale bar, 400 nm. Schematic depicting the hypothesized architecture of recombination factors at the crossover-designated site is shown on the right. (D) Representative SIM images of spread gonads from the region 2 in the diagram above and a segment of SC stretch from control and CDK-2-depleted animals. MSH-5 (green) and SYP-2 (red) staining are shown. Blue arrowhead in control indicates the SC bubble at the crossover site. Scale bar, 2 μm.

Examination of spread nuclei using 3D-SIM further revealed a failure to establish normal crossover site architecture in CDK-2-depleted germlines. Recent work has shown that crossover-designated sites display a distinct spatial organization of recombination factors (Jagut et al., 2016; Woglar and Villeneuve, 2018). Specifically, cohorts of MSH-5 and the Bloom helicase HIM-6 are each detected as orthogonally localized doublets. This orientation is interpreted to reflect their association with different parts of an underlying dHJ, with COSA-1 localizing at the center of the cross formed by the HIM-6 and MSH-5 doublets (Woglar and Villeneuve, 2018). Whereas this organization was detected in late pachytene in controls, MSH-5 and HIM-6 were not detected as doublets in CDK-2- depleted germ cells **(Figure 3C)**, consistent with a failure to form crossover-specific intermediates. Additionally, bubble-like SC structures have been detected at crossover-designated sites and are proposed to promote crossover maturation by encapsulating and protecting crossover-designated sites from anti-crossover activities (Woglar and Villeneuve, 2018). While SC bubbles surrounding MSH-5 foci were detected in controls, such structures were not found in CDK-2 depleted germlines **(Figure 3D)**. Taken together, our data support that CDK-2 is required for maturation of early recombination sites into crossover-specific recombination intermediates.

### Evidence for *in vivo* phosphorylation of MSH-5 by CDK-2

Experiments in which we co-expressed recombinant CDK-2-6xHis and GST- COSA-1 in insect cells provided evidence for the ability of CDK-2 to form a complex with COSA-1 *in vitro*. Although both proteins were largely insoluble despite optimization, we were able to purify CDK-2-6xHis from the soluble fraction using Ni-NTA beads and found that GST-COSA-1 was co-purified with CDK-2-6xHis at an apparent stoichiometric ratio **(Figures S3A-B)**.

Given the requirement for CDK-2/COSA-1 in crossover designation, we hypothesized that CDK-2/COSA-1 might phosphorylate pro-crossover factors, such as the MutSγ complex and the ZHP proteins, to modulate their functions. We focused on MSH-5, as it has been previously demonstrated to be a CDK substrate *in vitro* (Yokoo et al., 2012) and contains numerous CDK consensus motifs ([S/T]P) in its highly disordered C-terminal tail **(Figure 4A)**. Using mass spectrometry, we mapped three sites on the MSH-5 C-terminal tail (T1009, T1109, and S1278) that are phosphorylated by human CDK1 *in vitro*. To determine whether MSH-5 is indeed phosphorylated by CDK-2/COSA- 1 *in vivo*, we attempted to raise phospho-specific antibodies against these three sites and successfully generated an antibody that specifically detects MSH-5 phosphorylated at T1009 (MSH-5 pT1009). Immunofluorescence of spread nuclei revealed that the MSH-5 pT1009 signal was found at numerous recombination sites in early pachytene and became enriched at crossover-designated sites in late pachytene **(Figures S3C and 4B**), in alignment with the localization of MSH-5 and CDK-2/COSA-1 during meiotic progression. Further, the recognition of phosphorylated MSH-5 was eliminated in *msh-5* worms carrying the T1009A mutation **(Figure S3D)**, demonstrating the specificity of our antibody. Importantly, the signal for MSH-5 pT1009 was abolished from recombination sites in CDK-2-depleted gonads **(Figure 4B)**, indicating that the MSH-5 C-terminal tail is phosphorylated in a CDK-2-dependent manner *in vivo*.

**Figure 4.**
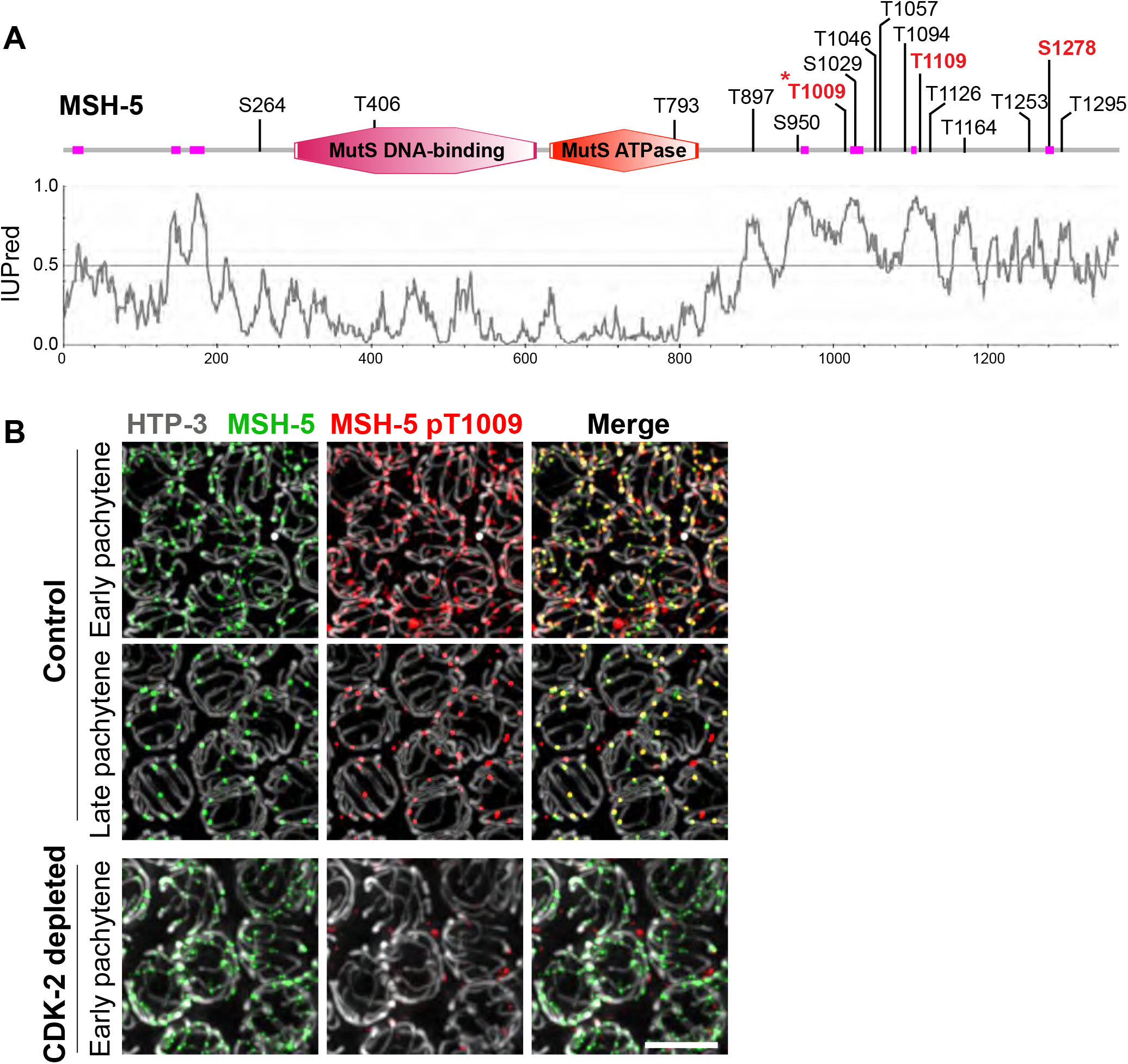
CDK-2 is responsible for MSH-5 phosphorylation within its C-terminal tail. (A) Schematic showing the domain structure and putative CDK phosphorylation sites in MSH-5 (adopted from the SMART database). Low-complexity regions are shown in magenta, and *in vitro* phosphorylation sites mapped by mass spectrometry analysis are indicated in red. An asterisk indicates the MSH-5 pT1009 epitope. The IUPred disorder score for MSH-5 is shown below. (B) Immunofluorescence images of spread pachytene nuclei from control (no auxin) and CDK-2-depleted germline (1 mM auxin treatment for 24 hr). Non-specific signals from the MSH-5 pT1009 antibody that do not colocalize with MSH-5 were occasionally detected at the nuclear periphery. Staining for HTP-3 (white), MSH-5 (green), and MSH-5 pT1009 (red) is shown. Scale bar, 5 μm.

### The C-terminal tail of MSH-5 is essential for accumulation of pro-crossover factors at recombination sites

Whereas the N-terminal 60% of the MSH-5 protein shows a high level of conservation with its orthologs throughout the eukaryotic kingdoms, the long C-terminal tail is unique to its Caenorhabditis orthologs (**Figure S4A**). Moreover, primary sequence conservation within this tail domain is very low among the Caenorhabditis MSH-5 orthologs, which are similar in that they contain multiple (8-23) CDK consensus motifs embedded within a protein segment predicted to be highly disordered (**Figure S4B**). These features suggest that the presence of the disordered tail and/or its ability to serve as a substrate for CDK-2 might contribute to the essential functions of MSH-5 in meiosis.

To address this, we used CRISPR to generate worm strains expressing a series of MSH-5 C-terminal truncations by inserting V5 coding sequences followed by premature stop codons (**Figure 5A**). Western blot analysis showed that all four truncated proteins (Δ178 aa, Δ270 aa, Δ339 aa, and Δ414 aa) are expressed at their expected sizes, albeit at reduced levels **(Figure S5A)**. Strikingly, truncations of MSH-5 led to a marked reduction in egg viability, and the percent males among surviving progeny also increased, reflecting meiotic impairment **(Figure S5B)**. In particular, animals harboring two large truncations (*msh-5::V5^Δ339 aa^* and *msh-5::V5^Δ414 aa^*) displayed an average of ∼11 DAPI bodies in diakinesis oocytes **(Figures 5B-C and S5C)** and only 0-1 bright COSA-1 foci in late pachytene nuclei **(Figures 5D and S5D)**, indicating that the C-terminal tail of MSH-5 is crucial for its function in crossover formation. Immunofluorescence of spread gonads from *msh-5^Δ339 aa^* mutants revealed that COSA-1 was diffusely present in the nucleoplasm **(Figure 5E)**. Additionally, after spreading, dim COSA-1 foci were detected at up to 6 sites along chromosome axes, which colocalized with ZHP-3 and the truncated MSH-5 (**Figures 5E-F**). However, ZHP-3 signal persisted along the SC in *msh-5^Δ339 aa^* mutants **(Figure 5F)**, and most recombination intermediates failed to mature into crossovers. Thus, the C-terminal tail of MSH-5 is essential for accumulation of pro-crossover factors at COSA-1-marked recombination sites.

**Figure 5.**
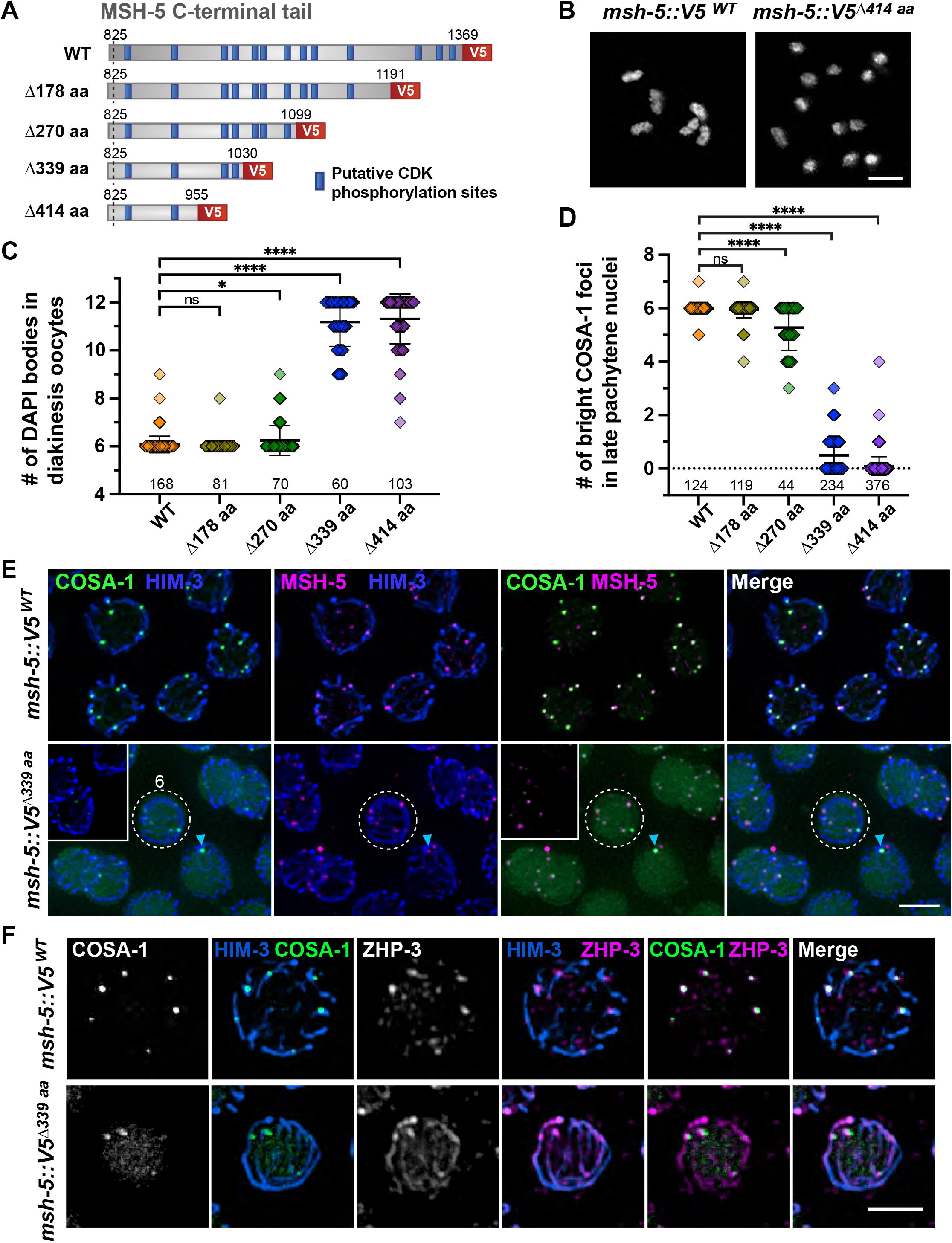
The disordered C-terminal tail of MSH-5 is essential for accumulation of pro-crossover factors at recombination sites. (A) Diagram of MSH-5 C-terminal truncation alleles. (B) DAPI-stained chromosomes in diakinesis oocyte nuclei from *msh-5::V5^WT^* and *msh-5::V5^Δ414aa^* animals. Scale bar, 3 μm. (C) Quantification of the number of DAPI bodies in diakinesis oocytes from indicated genotypes. Numbers of nuclei scored are shown below the graphs. Mean ± SD is shown. ****, p < 0.0001; *, p = 0.0128; ns, not significant by Mann-Whitney test. (D) Quantification of the number of bright COSA-1 foci per nucleus from indicated genotypes. Numbers of nuclei scored are shown below. Mean ± SD is shown. ****, p <0.0001; ns, not significant by Mann-Whitney test. (E) Spread gonads from *msh-5::V5^WT^* and *msh-5::V5^Δ339aa^* animals were stained for GFP::COSA-1 (green), HIM-3 (blue), and MSH-5 (magenta). The images from *msh-5::V5^Δ339aa^* were overexposed to visualize dim COSA-1 foci; insets represent settings comparable to the wild-type images. Late pachytene regions are shown. A representative nucleus with six faint COSA-1 foci in the *msh-5^Δ414aa^* mutant is highlighted by a white dotted circle. A bright COSA-1 focus that colocalizes with MSH-5 in the *msh- 5^Δ339aa^* is indicated by a cyan arrowhead. Scale bar, 3 μm. (F) Immunofluorescence images of spread late pachytene nuclei from *msh-5::V5^WT^* and *msh-5::V5^Δ339aa^* animals showing GFP::COSA-1 (green), HIM-3 (blue), and ZHP-3 (magenta). Scale bar, 3 μm.

### Phosphosites within the MSH-5 C-terminal tail contribute to the pro-crossover activity of the MutSγ complex

To determine the significance of phosphorylation of the MSH-5 tail, we sequentially mutated the codons corresponding to the 13 predicted and/or mapped CDK phosphorylation sites in its C-terminus using CRISPR **(Figures 6A and S6A)**. Contrary to our initial expectations based on the truncation mutants, animals homozygous for *msh*- 5 phospho-mutations (4A, 11A, and 13A) did not show obvious phenotypes in meiosis, exhibiting nearly normal progeny viability **(Figure S6B)**. Even the *msh-5::V5^13A^* mutant, in which all 13 CDK consensus sites were mutated to alanines, was able to designate crossovers and form bivalents as in wild type **(Figures S6C-E and 6B)**.

**Figure 6.**
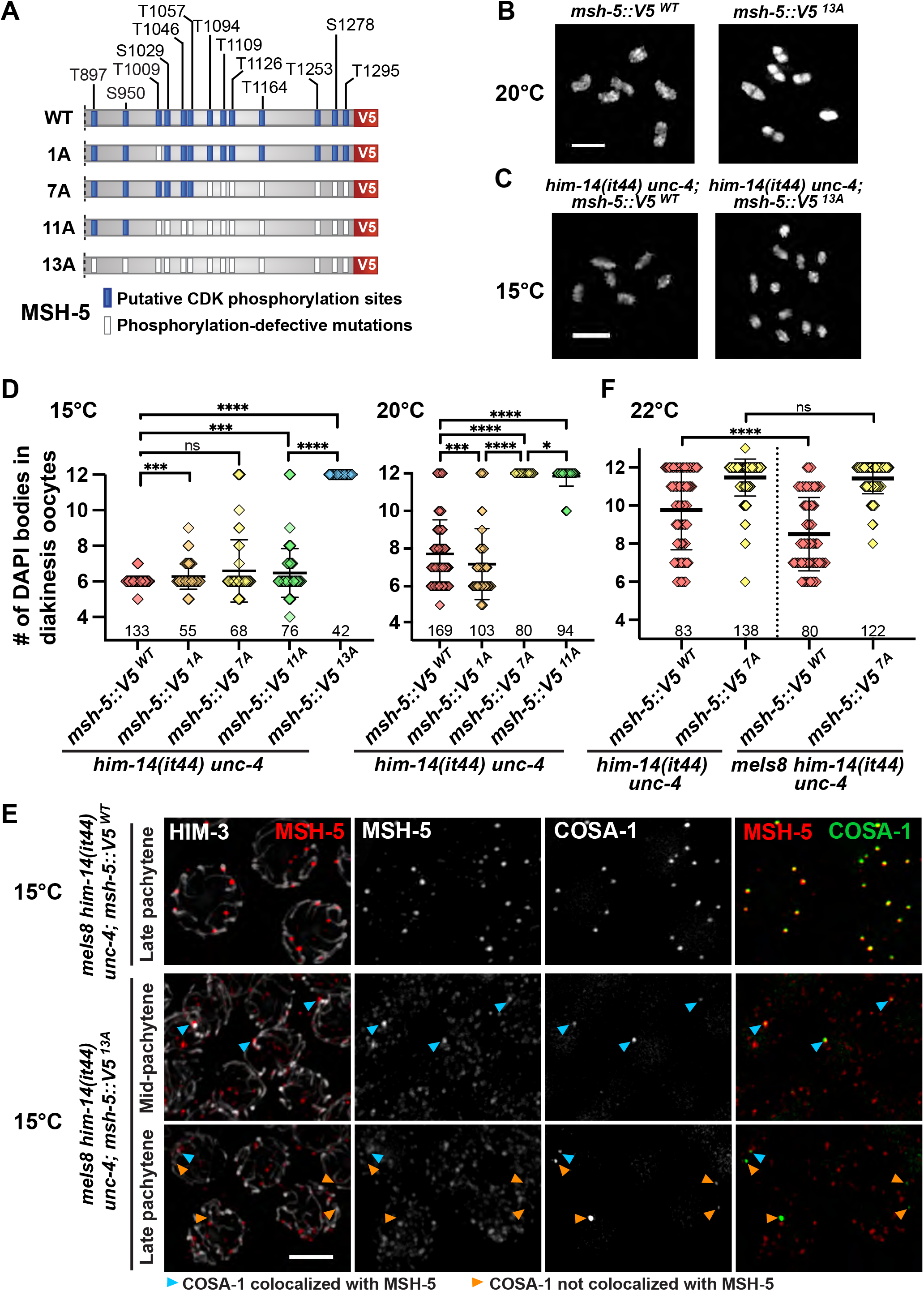
Phosphosites within the C-terminal tail of MSH-5 contributes to the pro- crossover activity of the MutSγ complex. (A) Diagram of *msh-5* mutant series harboring phosphorylation-defective mutations within its C-terminal tail, indicating with white boxes the positions of S/T residues that were replaced by A residues. (B) Oocyte nuclei at diakinesis from *msh-5::V5^WT^* and *msh- 5::V5^13A^*. Adult hermaphrodites grown at 20°C (48 hrs post L4) were stained with DAPI. Scale bar, 4 μm. (C) Oocyte nuclei at diakinesis from *him-14(it44) unc-4; msh-5::V5^WT^* and *him-14(it44) unc-4; msh-5::V5^13A^*. L4 hermaphrodites were grown at 15°C for 48 hr and were dissected for DAPI staining. Scale bar, 4 μm. (D) Graph showing the number of DAPI bodies in diakinesis oocytes from indicated genotypes grown at 15°C (left) and 20°C (right). Numbers of nuclei scored are shown below. Mean ± SD is shown. ****, p<0.0001; ***, p=0.0004-0.0009; *, p=0.0158; ns, not significant by Mann-Whitney test. (E) Immunofluorescence images of spread pachytene nuclei from indicated genotypes grown at 15°C showing staining for HIM-3 (white), MSH-5::V5 (red), and GFP::COSA-1 (green). Cyan arrowheads indicate COSA-1 foci that colocalize with MSH-5, while orange arrowheads indicate the ones that do not overlap with MSH-5. Scale bar, 3 μm. (F) Graph showing the number of DAPI bodies in diakinesis oocytes from indicated genotypes grown at 22°C. Numbers of nuclei scored are shown below. Mean ± SD is shown. ****, p<0.0001; ns, not significant by Mann-Whitney test.

As the conserved presence of multiple CDK phosphorylation sites suggests a functional importance, we hypothesized that preventing phosphorylation of the MSH-5 C-terminus did not cause overt phenotypes because phosphorylation is only one of multiple parallel pathways that converge to ensure a robust outcome of meiosis. Thus, we utilized a temperature-sensitive allele of *him-14/msh-4,* which encodes the heterodimeric partner of MSH-5 in the MutSγ complex, as a sensitized genetic background to reveal a functional deficit for *msh-5* phosphomutants. *him-14*(*it44)* harbors a missense mutation (D406N) within its conserved DNA-binding domain and is characterized by a temperature-sensitive reduction in crossover formation (Zalevsky et al., 1999; Zetka and Rose, 1995). Analysis of phosphosite mutations in this sensitized context provided clear evidence that phosphorylation of the MSH-5 tail functions to augment the pro-crossover activity of MutSγ in vivo.

First, we found that the *msh-5::V5^13A^* mutation showed strong synthetic phenotypes with *him-14(it44)* at 15°C, a temperature that is normally permissive for *him- 14(it44)* **(Figures 6C and 6D)**. Whereas DAPI staining of diakinesis oocytes did not reveal meiotic defects in *him-14(it44); msh-5::V5^WT^* control animals at 15°C, 12 univalents were consistently observed in *him-14(it44); msh-5::V5^13A^* oocytes, indicating a failure of crossover formation. Further, nuclear spreading and immunofluorescence revealed that while wild-type MSH-5 pared down to colocalize with six bright COSA-1 foci at crossover-designated sites in *him-14(it44)* at 15°C, the MSH-5^13A^ protein persisted as numerous foci throughout pachytene in the *him-14(it44); msh-5::V5^13A^* double mutant **(Figure 6E)**, similar to the phenotypes observed in CDK-2-depleted germlines **(Figure 3)**. Moreover, in *him-14(it44); msh-5::V5^13A^* animals, 1-3 COSA-1 foci initially associated with MSH-5, but this colocalization was lost in late pachytene (only 33% of COSA-1 signals colocalized with MSH-5 in *him-14(it44); msh-5::V5^13A^* vs. 98% in *him-14(it44); msh-5::V5^WT^*) **(Figures 6E and S7A)**. Interestingly, ZHP-3 was detected together with COSA-1 in late pachytene nuclei of *him-14(it44); msh-5::V5^13A^* **(Figure S7B)**, suggesting that phosphorylation of the MSH-5 C-terminus is required for retaining the association of MutSγ with other pro-crossover factors in *him-14(it44)* animals.

Second, analysis of *him-14(it44)* double mutant animals carrying phospho-null mutations at 1, 7, or 11 phosphosites (1A, 7A, 11A) further revealed that multiple phosphorylation sites in the MSH-5 tail can contribute to promoting MutSγ activity **(Figures 6A, 6D, and S7C)**. The *him-14(it44); msh-5::V5^11A^* mutant was particularly informative, as analysis of diakinesis oocytes revealed only mild meiotic impairment at 15°C, in striking contrast to the complete failure of crossover formation observed in *him- 14(it44); msh-5::V5^13A^* oocytes under the same conditions. This suggests that phosphorylation of as few as two sites within the MSH-5 tail can sustain sufficient pro-crossover activity of the partially compromised MutSγ complexes. However, severe impairment of crossover formation is observed in both *him-14(it44); msh-5::V5^11A^* and *him-14(it44); msh-5::V5^7A^* when MutSγ function is further compromised at the semi-permissive temperature of 20°C **(Figure 6D right).** This suggests that phosphorylation at additional sites can contribute to augmentation of MutSγ activity.

### MSH-5 phosphosites are required for suppression of the *him-14(it44)* crossover deficit by elevated COSA-1 levels

Overexpression of COSA-1 by the *meIs8* transgene can partially alleviate the crossover deficit observed at semi-permissive temperatures in the *him-14(it44)* mutant (Girard et al., 2021). This suppression is hypothesized to reflect elevated CDK-2/COSA- 1 activity, which compensates for the impaired MutSγ activity in *him-14(it44)* through hyper-phosphorylation of downstream targets, including MSH-5. To test whether suppression of *him-14(it44)* by *meIs8* was dependent on phosphosites in the MSH-5 C-terminal tail, we compared the effect of *meIs8* in *him-14(it44); msh-5::V5^WT^* and *him- 14(it44); msh-5::V5^7A^* genetic backgrounds. Experiments were conducted at 22°C, a semi-permissive temperature at which the suppression of *him-14(it44)* by *meIs8* is evident. At 22°C, *him-14(it44); msh-5::V5^WT^* diakinesis oocytes displayed an average of 9.8 DAPI bodies, reflecting a mixture of bivalents and univalents resulting from a partial impairment of crossover formation, but the number of DAPI bodies was reduced to 8.5 in the presence of the *meIs8* transgene **(Figures 6F and S7D),** indicating an increase in bivalent formation reflecting increased success in crossover formation. However, there was no significant difference in the number of DAPI bodies, with or without the *meIs8* transgene, in the *him-14(it44); msh-5::V5^7A^* background **(Figures 6F and S7D)**. This finding strongly suggests that phosphorylation of the MSH-5 C-terminal tail is required for suppression of *him-14(it44)* phenotypes by the elevated CDK-2/COSA-1 activity, supporting the conclusion that the MSH-5 tail is the major CDK-2/COSA-1 target whose phosphorylation is responsible for augmenting the residual MutSγ activity in *him-14(it44)* animals.

## DISCUSSION

### CDK-2 partners with COSA-1 to form a dedicated meiotic CDK/cyclin complex

Despite the long-standing evidence that mammalian CDK2 localizes to interstitial sites on chromosomes during meiosis (Ashley et al., 2001), it has been challenging to elucidate its function in crossover recombination. Here we demonstrate that *C. elegans* CDK-2 localizes to recombination intermediates and partners with COSA-1 to promote crossover formation. The CDK-2/COSA-1 complex functions to convert a subset of meiotic DSBs into interhomolog crossovers by stabilizing crossover-specific recombination intermediates. As CDK-2 is dispensable for homolog pairing and synapsis in *C. elegans*, our work unequivocally establishes a role for CDK-2 in crossover designation.

### CDK-2/COSA-1 promotes crossover designation through phosphorylation of MSH- 5

Here we identify MutSγ as a key meiotic target of CDK-2/COSA-1 and demonstrate that MSH-5 is phosphorylated in a CDK-2-dependent manner within its disordered C-terminal tail. As mutating all 13 C-terminal CDK motifs in MSH-5 does not result in meiotic defects, CDK-2 likely has additional substrates essential for crossover formation. However, severe consequences of phosphosite loss are evident when the activity of MutSγ is compromised, indicating the importance of phosphorylation within the MSH-5 tail for enabling success of meiosis under suboptimal conditions. The significance of this kinase-substrate relationship is further supported by dosage suppression, in which extra copies of the *cosa-1* gene enable *him-14(it44)* animals to form higher levels of crossovers at a semi-permissive temperature (Girard et al., 2021). We have shown that suppression of *him-14(it44)* phenotypes by COSA-1 overexpression requires MSH-5 phosphorylation within its C-terminal region. Thus, MSH-5 is a key substrate of CDK-2/COSA-1, and phosphorylation within the MSH-5 C-terminal tail potentiates the overall activity of MutSγ in stabilizing crossover-specific recombination intermediates **(Figure 7)**.

**Figure 7.**
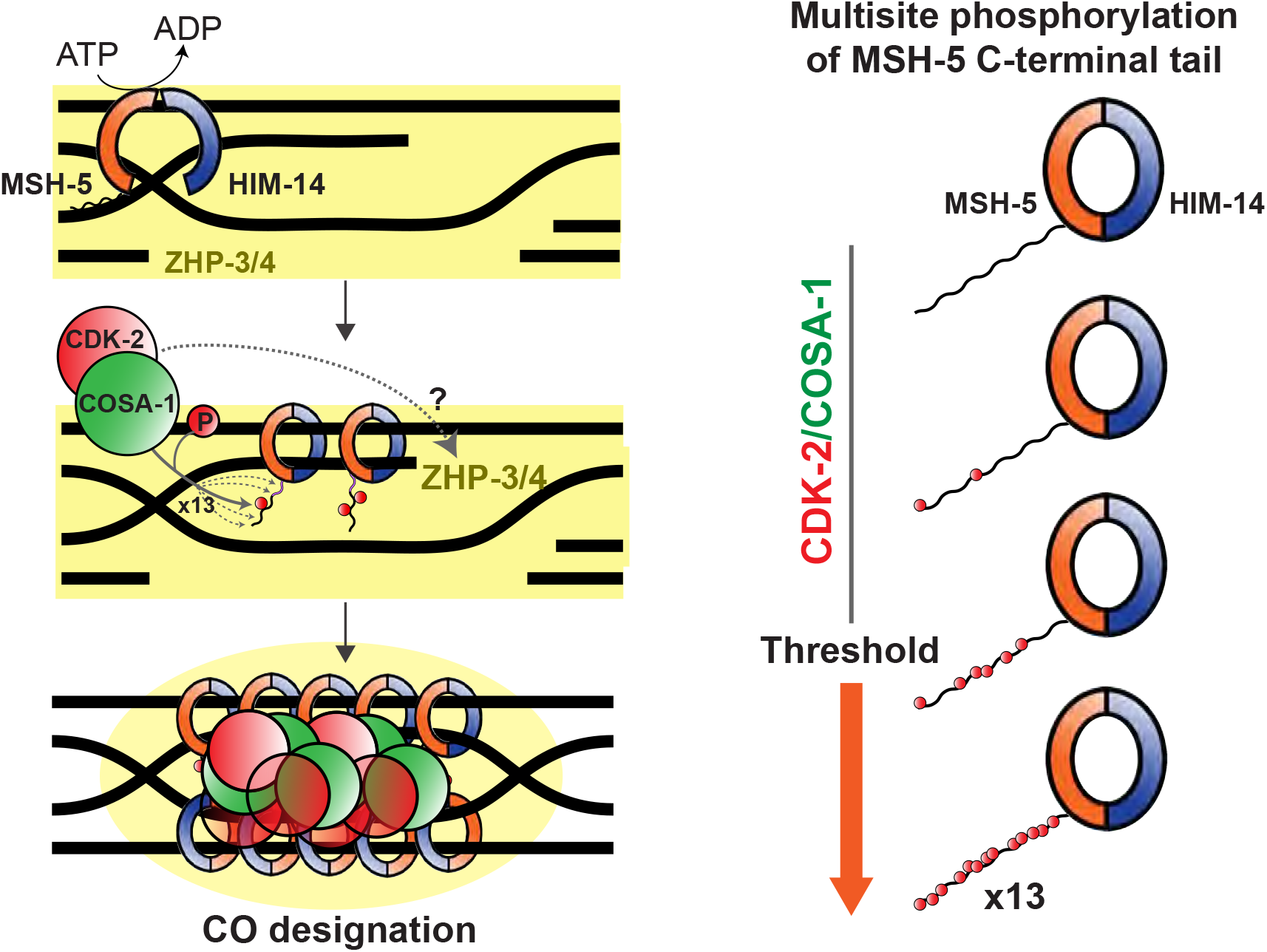
Model for robust designation of meiotic crossover sites through CDK- 2/COSA-1-dependent multisite phosphorylation of the MutSγ complex, positive feedback, and scaffold-like properties of the MSH-5 tail. The MutS**γ** complex recognizes and binds nascent recombination intermediates. CDK- 2/COSA-1 is also recruited to the early recombination sites and phosphorylates MSH-5 in its disordered C-terminal tail. As the recombination intermediate matures and recruits more MutSγ and CDK-2/COSA-1, more sites in the MSH-5 tail are phosphorylated. CDK- 2/COSA-1 likely phosphorylates other substrates essential for crossover formation. Multisite phosphorylation of the MSH-5 C-terminal tail generates an ultrasensitive response that potentiates the overall activity of MutSγ to stabilize crossover-specific recombination intermediates. Phosphorylation within the MSH-5 tail also helps retain CDK-2/COSA-1 and ZHP-3/4 (depicted in yellow), thereby providing positive feedback and conferring a switch-like behavior that contributes to the robustness of crossover designation.

### The C-terminal tail of MSH-5 as a scaffold to accumulate other pro-crossover factors

CDK phosphorylation sites in MSH-5 are clustered within its disordered C-terminal domain, which we have shown to be essential for crossover formation. Since the MSH-5 C-terminal tail is outside of its enzymatic core or the dimerization interface mapped for human MSH4 and MSH5 (Snowden et al., 2008), it is unlikely that deleting the C-terminal tail affects the ATP hydrolysis rate or the formation of the HIM-14/MSH-5 heterodimer. In worms expressing a truncated MSH-5 (*msh-5^Δ339 aa^*), MSH-5 localizes to no more than six COSA-1-marked recombination sites, suggesting that crossover site designation may have occurred. However, pro-crossover factors do not accumulate to wild-type levels at these sites, depletion of ZHP-3 from along the length of the SC does not occur, and lack of chiasmata connecting homologs at diakinesis indicates a failure to process these recombination intermediates into crossovers **(Figure 5)**. We propose that the C-terminal tail of MSH-5 serves as a scaffold to accumulate proteins required for crossover formation. Indeed, intrinsically disordered proteins frequently contain short linear motifs that mediate interactions with diverse targets and have emerged as major hubs in cellular signaling (Wright and Dyson, 2015). Recent work in mice has identified a novel proline-rich protein, PRR19, that functions with CNTD1 to promote crossover formation (Bondarieva et al., 2020). Intriguingly, the MSH-5 tail is also enriched in prolines (CDK is a proline-directed kinase) and thus may act as a functional substitute for PRR19 to stably recruit CDK-2/COSA-1 in *C. elegans*. Determining how the MSH-5 tail mediates higher-order assemblies of pro-crossover factors will be an important topic for future research.

The phenotype observed in *msh-5^Δ339 aa^* animals is in sharp contrast to that in *msh- 5::V5^13A^; him-14(it44)*, where 1-3 bright foci of COSA-1 initially localize to a subset of MutSγ-positive recombination intermediates but lose their association in late pachytene **(Figure 6E)**. Thus, although the MSH-5 C-terminal tail itself is required for concentrating pro-crossover factors at recombination sites, it appears to have deleterious effects on their retention in its unphosphorylated form. We speculate that the C-terminal tail of MSH- 5 may also act as a scaffold for recruiting anti-recombinases, which is counteracted by CDK-2-dependent phosphorylation. Recent evidence in *S. cerevisiae* has demonstrated that phosphorylation of the N-terminal degron within Msh4 protects it from proteolysis at recombination sites, thereby activating its pro-crossover activity (He et al., 2020). However, phosphorylation of the MSH-5 C-terminal tail does not seem to have similar stabilizing effects as we did not observe an increased level of MSH-5 in our C-terminal truncation or phosphosite mutants (data not shown). As ZHP-3 is depleted from the SC in *msh-5::V5^13A^; him-14(it44)* mutants while colocalizing with COSA-1, CDK-2/COSA-1 may also phosphorylate ZHP-3/4 and trigger their relocation from the SC to crossover-designated sites **(Figure 7)**.

### Multisite phosphorylation, positive feedback, and propensity for aggregation provide a robust mechanism for crossover designation

CDK phosphorylation sites are often clustered in disordered regions (Holt et al., 2009), and multisite phosphorylation by CDK can set a threshold to elicit an ultrasensitive response (Ferrell and Ha, 2014; Nash et al., 2001). Our analysis of *msh-5* phosphosite mutants revealed that crossover formation becomes highly sensitive to the number of phosphorylation sites available in the MSH-5 C-terminal tail when the activity of MutSγ is compromised by a temperature-sensitive mutation in *him-14*. At more permissive temperatures where HIM-14 is largely functional, fewer phosphosites are needed to achieve a threshold level of MutSγ activity required to ensure crossover formation. Conversely, under more restrictive conditions where HIM-14 is less functional, more phosphosites are required to achieve a threshold level of MutSγ activity. Although all MSH-5 orthologs in the *Caenorhabditis* species possess numerous CDK consensus motifs in the disordered C-terminal region, these sites are poorly conserved. Thus, we speculate that the number of phosphorylation sites, rather than their exact position, influences the pro-crossover activity of MutSγ, in line with several precedents controlled by multisite phosphorylation (Doan et al., 2006; Strickfaden et al., 2007). Further, as phosphorylation within the MSH-5 tail promotes the stable association of CDK-2/COSA- 1 to recombination sites, it can also generate positive feedback that further enhances MSH-5 phosphorylation, thereby conferring a switch-like behavior that contributes to the robustness of crossover designation.

We further speculate that the propensity for MSH-5 to form aggregates in late pachytene germ cells, revealed upon CDK-2 depletion, may also be a feature that promotes robustness of the crossover designation mechanism. Given the intrinsically disordered protein sequence in which phosphosites are embedded, we hypothesize that the MSH-5 C-terminal tail may have a capacity to undergo phase separation that can be modulated by phosphorylation, which could potentially contribute to the formation of ellipsoidal protein structures that have long been recognized as “recombination nodules” (Carpenter, 1975). The idea that formation of biomolecular condensates may contribute to crossover designation is supported by recent modeling of cytological data from *Arabidopsis,* in which HEI10 is proposed to accumulate at few sites through diffusion-mediated coarsening at the expense of smaller foci (Morgan et al., 2021). By enriching pro-crossover factors at crossover-designated intermediates while depleting them from other recombination sites, the formation of biomolecular condensates can serve as a general mechanism for controlling both crossover designation and positioning.

## ACKNOWLEDGEMENTS

We thank Needhi Bhalla for the ZHP-3 antibody and Abby Dernburg for the HIM-8 antibody. We also thank Yana Li at the Eukaryotic Tissue Culture Facility at Johns Hopkins for baculovirus production in insect cells and members of the Yumi Kim and Anne Villeneuve labs for critical reading of this manuscript. N2 worms were provided by the CGC, which is funded by the NIH Office of Research Infrastructure Program (P40OD010440). JH is supported by the NIH Predoctoral Fellowship (F31HD100142). This work was supported by funding from the National Institutes of Health to SA (R01GM98200), AV (R01GM067268 and R35GM126964), and YK (R35GM124895).

## AUTHOR CONTRIBUTIONS

JH, AV, and YK conceived and designed the study. JH performed most experiments. AW and JH conducted the SIM imaging and imaging using phospho-specific antibodies. CA constructed the strains containing *him-14(it44),* and CA, JH, and CS scored DAPI-staining bodies in diakinesis. KK and VR assisted JH in the construction of worm strains and plasmids for protein expression. SYC and SA mapped CDK phosphorylation sites on MSH-5 *in vitro*. JH, AV, and YK wrote and revised the manuscript.

## MATERIALS AND METHODS

### *C. elegans* strains and egg count assay

All strains used in this study were maintained on NGM plates seeded with OP50- 1 bacteria under standard conditions as described in (Brenner, 1974). All experiments were performed at 20°C except where noted. All *C. elegans* strains were derived from Bristol N2 background. **Tables S1 and S3** summarize all mutations and strains used in this study. To determine egg viability, brood size, and male progeny, L4 hermaphrodites were picked onto individual plates and moved every 12 hours. Eggs were counted upon time of transfer, and surviving males and hermaphrodites were counted upon reaching adult stage.

### Generation of *C. elegans* strains by CRISPR-mediated genome editing

The strains expressing CDK-2::AID::3xFlag and variants of MSH-5::V5 were generated by Cas9/CRISPR-mediated homologous recombination (Dokshin et al., 2018). Worms expressing GFP::COSA-1 and TIR1::mRuby under the *sun-1* promoter were injected with 0.25 μg/μL of Cas9 (2 nmol) complexed with 10 μM tracrRNA/crRNA oligos (IDT), 40 ng/μL of pRF4::*rol-6*(*su1006*), and a ssDNA oligo (200 ng/μL) (IDT) with 35 bp homology arms on both sides or gBlock (50 ng/μL) as a repair template (**Table S2**). Injections were performed using an Eppendorf Femtojet microinjector and a Narishige micromanipulator on an Olympus IX51 inverted microscope. F1 progeny were lysed and genotyped by PCR to detect successful CRISPR edits (**Table S2**). The correct insertion was validated by sequencing.

To mutate putative phosphorylation sites within the MSH-5 C-terminal tail, serine-to-alanine or threonine-to-alanine mutations were sequentially introduced by CRISPR. The truncation mutant series of MSH-5::V5 was generated by inserting a V5 tag with a GS linker (GGATCG) followed by a premature stop codon at desired truncation sites. The correct mutagenesis was validated by sequencing, and the strains were outcrossed with N2 3 times prior to analyses.

### Auxin-mediated depletion of CDK-2

Auxin-mediated degradation of CDK-2 from the *C. elegans* germline was performed as previously described (Zhang et al., 2015). Briefly, auxin plates were prepared by diluting a 400 mM auxin solution (indole-3-acetic acid in ethanol) into NGM, cooled after autoclaving, to a final concentration of 1 mM. Plates were dried at room temperature and stored at 4°C protected from light for up to one week prior to use. Plates were spread with concentrated OP50-1 bacterial cultures and incubated overnight at 37°C. Age-matched young adult hermaphrodites were picked and left for indicated hours as noted in each Figure at 20°C prior to immunofluorescence.

### Scoring DAPI-staining bodies in diakinesis oocytes of *him-14(it44)* animals

All strains carrying *him-14(it44) unc-4(e120)* were maintained at 20°C with the *mnC1* balancer. Homozygote L4 worms (Unc, non-green) were picked and transferred to experimental temperatures (15°C, 20°C or 22°C) for approximately 44 hours. To score DAPI-staining bodies in diakinesis oocytes, worms were picked onto a slide with a minimal volume of M9. Excess liquid was wicked away, and animals were fixed in 15 µl of 95% ethanol. Once dry, ethanol was reapplied, and this process was repeated a total of three times. The slides were mounted using VECTASHIELD containing DAPI (Vector Laboratories H-1200-10) and sealed. Slides were stored at 4°C for no longer than 4 days before imaging using a standard fluorescent microscope. DAPI bodies in the nuclei of diakinesis oocytes in the -1 to -3 positions were counted.

### Mapping CDK phosphorylation sites on MSH-5 *in vitro*

The full-length cDNA of *C. elegans* MSH-5 was cloned into the pTrcHis Topo Vector (Thermo Fisher K4410-01) to produce recombinant MSH-5 tagged with 6XHis at its N-terminus. Clones were verified by sequencing for orientation and sequence integrity. Protein expression was induced in BL21(DE3) cells (Sigma 69450) at 20°C using 1 mM isopropyl β-D-1-thiogalactopyranoside (dioxane free) for 16 hr. Proteins were then purified using Ni-NTA agarose (Qiagen 30210). The amount of purified protein was estimated both by conducting BCA assays (Thermo Fisher 23225) and by comparing band intensities of purified protein samples with a dilution series of 2 mg/mL Bovine Serum Albumin standard (Thermo Fisher 23209) on a Coomassie-stained SDS-PAGE gel. *In vitro* kinase reactions using purified CDK1/CyclinA2 kinase (Promega V2961) were carried out in a 25 μl reaction containing 100 ng of 6His-MSH-5, 50 μM cold ATP (Sigma), and 5U of active CDK1/CyclinA2 enzyme in 25 mM Tris pH7.0, 0.1 mM EGTA, 10 mM magnesium acetate. Reactions were terminated after 1 hour at 30°C. The protein was then purified using Phos-tag Agarose Resins (Wako AG-501) to enrich for the phosphoproteins and submitted to the TAPLIN Mass Spectrometry Facility for identification of phosphorylated peptides.

### Phosphopeptide antibody production and affinity purification

A synthetic phospho-peptide (TAIHIP[pT]PIQMGEAC) corresponding to the *C. elegans* MSH-5 sequence flanking threonine 1009 was generated using standard methods. The phospho-peptide was conjugated to KLH (Keyhole limpet hemacyanin) and injected into rabbits (Covance). To affinity-purify polyclonal MSH-5 pT1009 antibodies, immune serum was first passed through SulfoLink coupling resins (Thermo Fisher 20401) coupled to a non-phospho-peptide (TAIHIPTPIQMGEAC). Flow-through was then bound and eluted from phospho-peptide-coupled resins. The specificity of the antibodies was verified by dot blot and immunofluorescence of worm strains carrying phosphorylation-defective mutations in *msh-5*.

### Immunofluorescence

Young adult hermaphrodites (24 hr post L4) were dissected in Egg buffer (25 mM HEPES pH 7.4, 118 mM NaCl, 48 mM KCl, 2 mM EDTA, 5 mM EGTA, 0.1% Tween-20, 15 mM NaN3) and briefly fixed in 1% formaldehyde or for 5 minutes in 1% PFA (ZHP-3 staining shown in Figures 6 and S5B). Gonads were flash-frozen in liquid nitrogen, freeze-cracked and fixed in -20°C methanol for less than 1 minute. Fixed gonads were rehydrated in PBST (PBS and 0.1% Tween 20) and blocked with Roche Blocking reagent (Sigma 11096176001) for 35 minutes at room temperature. Samples were incubated with primary antibodies overnight at 4°C at the indicated dilutions: mouse anti-FLAG (1:200,

Sigma F1804), GFP VHH (1:200, Chromotek gt-250), guinea pig anti-HTP-3 (1:500) (MacQueen et al., 2005), chicken anti-HIM-3 (1:500) (Hurlock et al., 2020), rabbit anti-phospho-HIM-8/ZIMs (1:1000) (Kim et al., 2015), rabbit anti-MSH-5 (1:10000, SDI/Novus 38750002), rat anti-RAD-51 (1:200) (Rosu et al., 2013), rabbit anti-MSH-5 pT1009 (1:200, this study), mouse anti-V5 (1:200, Invitrogen R960-25), rabbit anti-DSB-2 (1:5000) (Rosu et al., 2013), rabbit anti-SYP-1 (1:300) (Brandt et al., 2020), rabbit anti-SYP-2 (1:200) (Colaiacovo et al., 2003), rabbit anti-SYP-5 (1:500) (Hurlock et al., 2020), guinea-pig anti-ZHP-3 (1:100) (Bhalla et al., 2008), and rat HIM-8 (1:500) (Phillips et al., 2005). Slides were washed with PBST and incubated with secondary antibodies for 30 minutes at room temperature at a 1:200 dilution (Sigma or Invitrogen Alexa 488, Alexa 555 or Alexa 647). Slides were washed again in PBST, stained with DAPI, and mounted in ProLong Gold (Invitrogen P36930). Slides were cured for 1-3 days and imaged using a DeltaVision^TM^ Elite system (GE) equipped with an 100x oil-immersion, 1.4 NA objective and a sCMOS camera (PCO). 3D image stacks were collected at 0.2 µm intervals, processed by iterative deconvolution (enhanced ratio, 20 cycles), and projected using the SoftWoRx package. Composite images were assembled and colored using Adobe Photoshop.

Chromosome spreading under high (Figures 1B, 1C, 3A, 3B, 3D, 4B, 5E, 5F, 6E, S2E, S3C, S3D and S5E) and low salt (Figures 1D and 3C) conditions was performed as previously described (Woglar and Villeneuve, 2018). Young adult hermaphrodites were dissected in 20 μl dissecting solution (0.1% v/v Tween-20, Hank’s Balanced Salt Solution (Gibco 24020117) (high salt- 85% v/v, low salt- 10% v/v)) on an ethanol-washed 22x40 mm or 22x22 mm coverslip. Dissected gonads were spread across the cover slip using a pipette tip in 50 μl of spreading solution (2.5% w/v paraformaldehyde, 2% w/v sucrose, 0.32% v/v Lipsol, 0.04% w/v Sarcosyl). Coverslips were dried at room temperature overnight and washed in -20°C methanol for 20 minutes. After rehydrating in PBST (3 x 5 min), samples were processed for immunofluorescence using antibodies at the concentrations listed above.

### Three-dimensional structured illumination microscopy (3D-SIM)

Samples were imaged, deconvolved and 3D-SIM reconstructed as conducted in (Pattabiraman et al., 2017). Slides were prepared by chromosome spreading (described above) and imaged as 125 nm spaced Z-stacks using a DeltaVision OMX Blaze microscopy system (GE) equipped with a 100x NA 1.40 objective. SoftWoRx was used to 3D-reconstruct images and correct for registration. Images were projected for display using maximum intensity projection in Fiji or SoftWoRx, and individual color channel contrast and brightness were adjusted for figures in Fiji.

### Expression and purification of the CDK-2/COSA-1 complex

Synthetic gene fragments (IDT) encoding the full-length open reading frame (ORF) of *C. elegans* CDK-2 and COSA-1 were cloned into pFastBac1 (Gibco). For protein purification, COSA-1 was tagged at its N-terminus with glutathione S-transferase (GST) and CDK-2 was C-terminally tagged with 6xHis. GST-COSA-1 and CDK-2-6xHis were co-expressed and purified from Sf9 cells using the Bac-to-Bac system (Life Technologies). Infected High-Five cells (Life Technologies) were lysed in 50 mM Tris pH 7.5, 20 mM imidazole, 500 mM NaCl, 0.5 mM EGTA, 1% NP-40, 1 mM DTT, 1 mM PMSF,

Complete Protease inhibitor Cocktail (Roche 11873580001) by dounce homogenization followed by sonication. 6xHis-tagged CDK-2 was purified using Ni-NTA resins (Qiagen 30210) and eluted in PBS, 500 mM imidazole, and 1 mM DTT. Purified proteins were separated by SDS-PAGE and analyzed by Coomassie blue staining and immunoblots.

### Immunoblots

Protein samples were separated by SDS-PAGE and transferred onto nitrocellulose membranes (GE Healthcare) using the Trans-Blot Turbo System (BioRad). Membranes were blocked in 5% milk in PBST for 1 hour at room temperature and incubated overnight at 4°C with agitation in primary antibody at the following dilutions: rabbit anti-His (1:1,000, Cell Signaling, 2365), rabbit anti-GST (1:1,000, Invitrogen, PA1-982A), mouse anti-V5 (1:1,000, Abcam, ab27671), and rabbit anti-SYP-2 (1:1000, (Colaiacovo et al., 2003)).

## SUPPLEMENTAL FIGURE LEGENDS

**Figure S1 (related to Figure 1).**
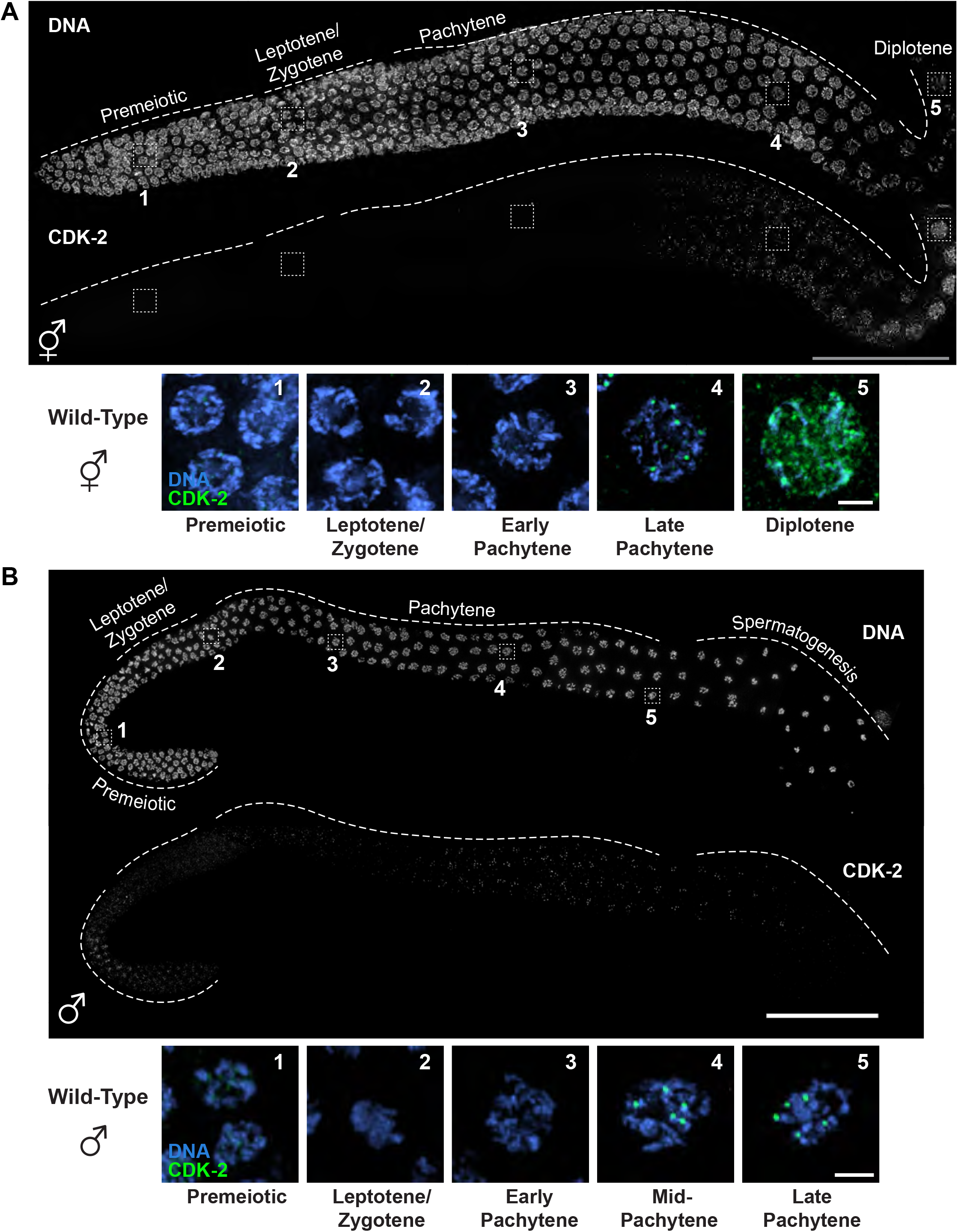
CDK-2 localizes to crossover sites in both hermaphrodite and male germlines. (A) Composite immunofluorescence image of a whole mount gonad dissected from an adult hermaphrodite expressing CDK-2::AID::3xFlag. DNA and Flag staining are shown. Scale bar, 50 μm. Below: Representative nuclei from the boxed regions stained for DNA (blue) and CDK-2 (green). Scale bar, 2 μm. (B) Composite immunofluorescence image of a whole mount gonad dissected from an adult male expressing CDK-2::AID::3xFlag. Scale bar, 50 μm. Below: Representative nuclei from the boxed regions stained for DNA (blue) and CDK-2 (green). Scale bar, 2 μm.

**Figure S2 (related to Figure 2).**
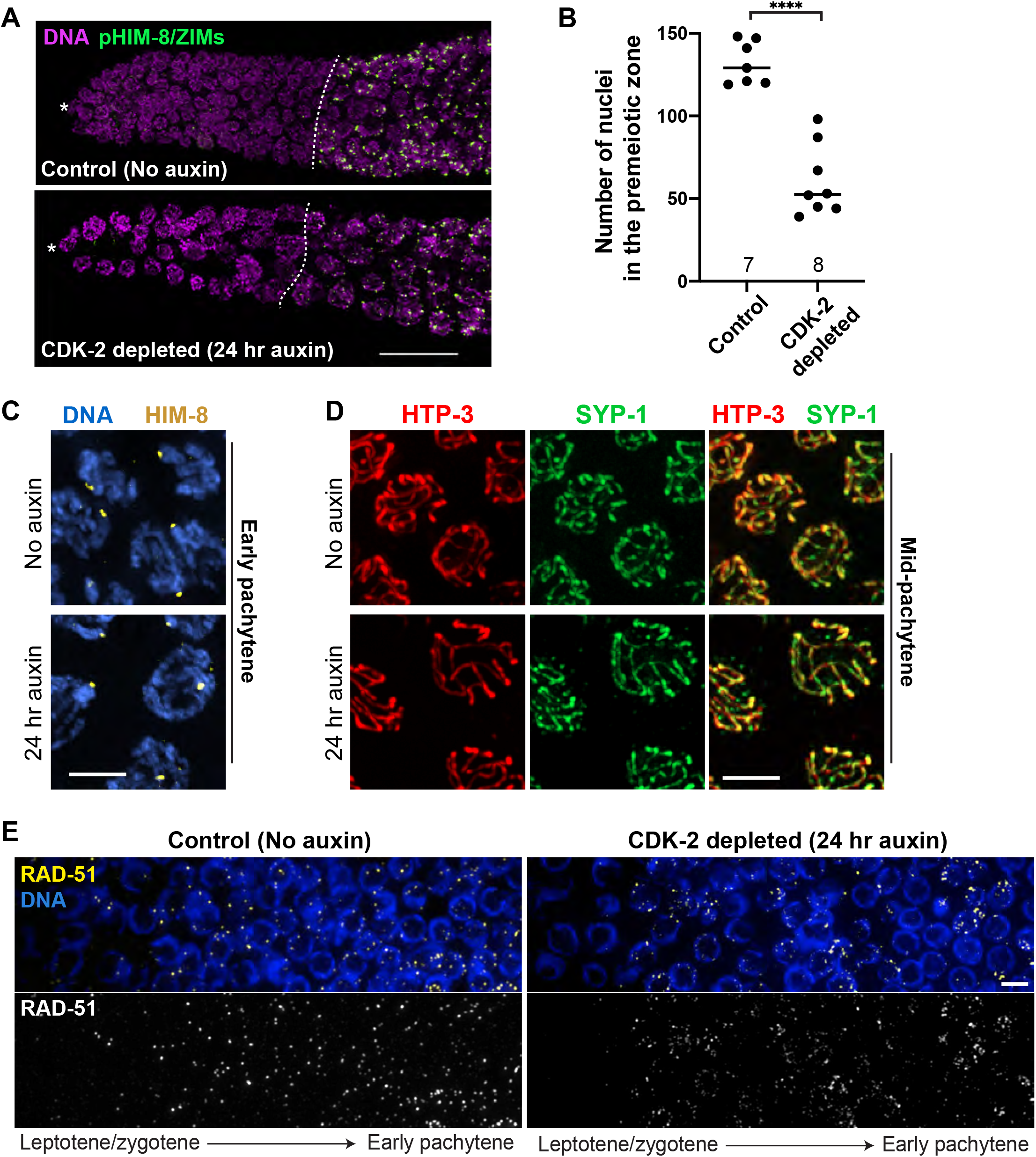
CDK-2 is dispensable for homolog pairing, synapsis, and initiation of meiotic recombination. (A) Young adult hermaphrodites expressing CDK-2::AID::3xFlag, GFP::COSA-1, and *P_sun-1_*::TIR1::mRuby were treated with or without 1 mM auxin for 24 hr. Composite immunofluorescence images of distal germlines from control (no auxin) and CDK-2- depleted (24 hr auxin) animals showing DNA (magenta) and pHIM-8/ZIMs staining (green) to mark meiotic onset. Asterisks indicate the distal tip of the germline, and dotted lines indicate meiotic entry. Scale bar, 15 μm. (B) Graph showing the number of nuclei in the premeiotic zone in control (n = 7) and CDK-2-depleted germlines (n = 8). ****p < 0.0001 by nonparametric unpaired t-test. (C) Immunofluorescence images of early pachytene nuclei from control (no auxin) and CDK-2-depleted (24 hr auxin) gonads stained for DNA (blue) and HIM-8 (yellow). Scale bar, 3 μm. (D) Immunofluorescence images of mid-pachytene nuclei from control (no auxin) and CDK-2-depleted (24 hr auxin) gonads stained for HTP-3 (red) and SYP-1 (green). Scale bar, 3 μm. (E) Immunofluorescence images of nuclei from leptotene/zygotene to early pachytene showing DNA (blue) and RAD-51 staining (yellow or white) in control (no auxin) and CDK- 2-depleted (24 hr auxin) gonads. Scale bar, 5 μm.

**Figure S3 (related to Figure 4).**
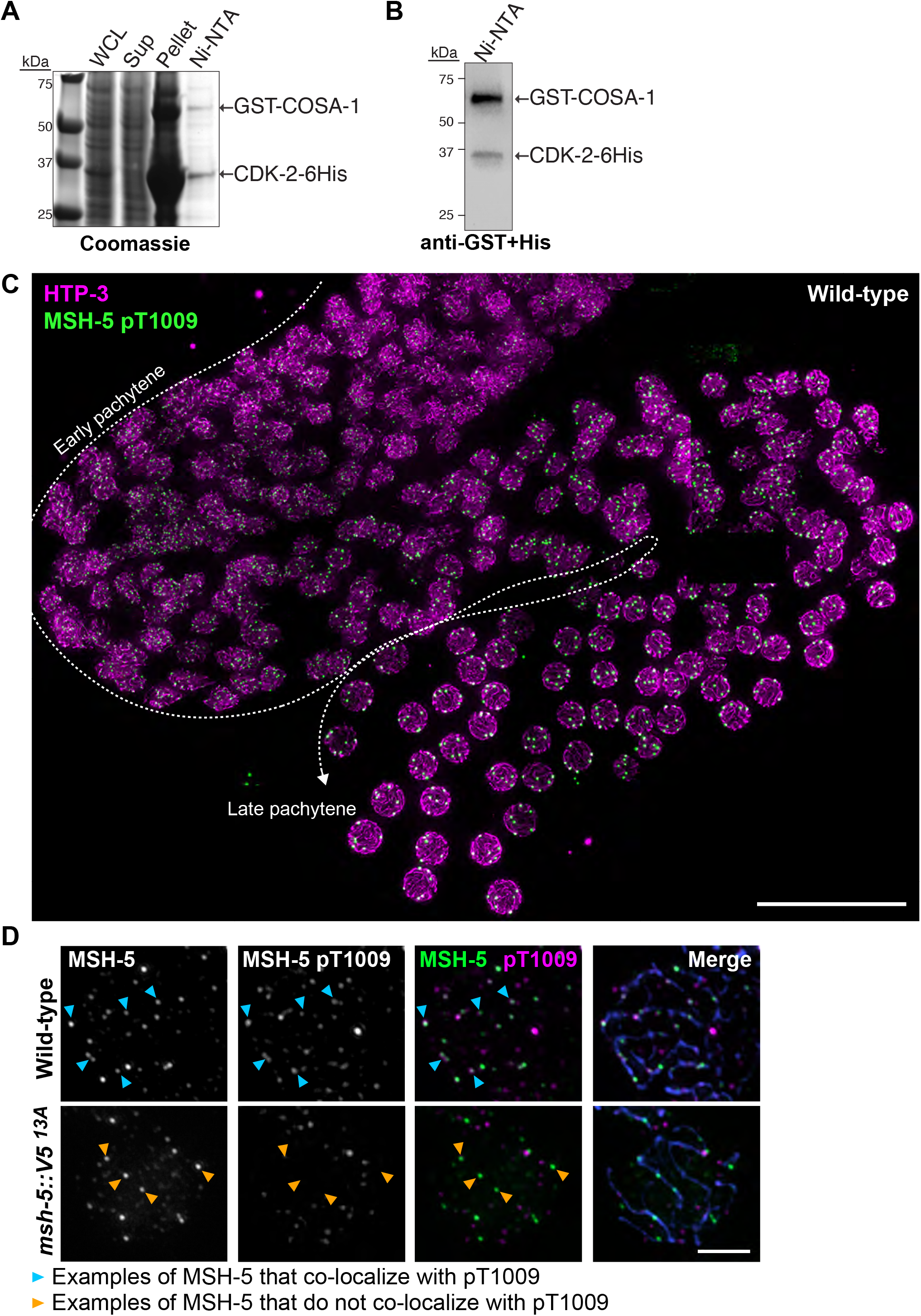
Purification of the CDK-2/COSA-1 complex and characterization of the MSH-5 pT1009 antibody. (A) Coomassie-stained SDS-PAGE gel showing the co-purification of CDK-2-6His and GST-COSA-1 using Ni-NTA beads. WCL: whole cell lysates; Sup: supernatant. (B) Immunoblot showing the purified GST-COSA-1/CDK-2-6His complex using both GST and His antibodies. (C) Composite immunofluorescence image of a spread wild-type gonad showing HTP-3 (magenta) and MSH-5 pT1009 (green). The direction of meiotic progression is indicated by the white dotted arrow. Scale bar, 15 μm. (D) Immunofluorescence images of late pachytene nuclei from wild-type and *msh-5::V5^13A^* mutant showing MSH-5 (green), MSH-5 pT1009 (magenta), and HIM-3 (blue). Scale bar, 5 μm. Examples of MSH-5 foci that colocalize with the MSH-5 pT1009 signal are indicated by cyan arrowheads, and MSH-5 foci that do not colocalize with pT1009 antibody signals are indicated by orange arrowheads.

**Figure S4 (related to Figure 5).**
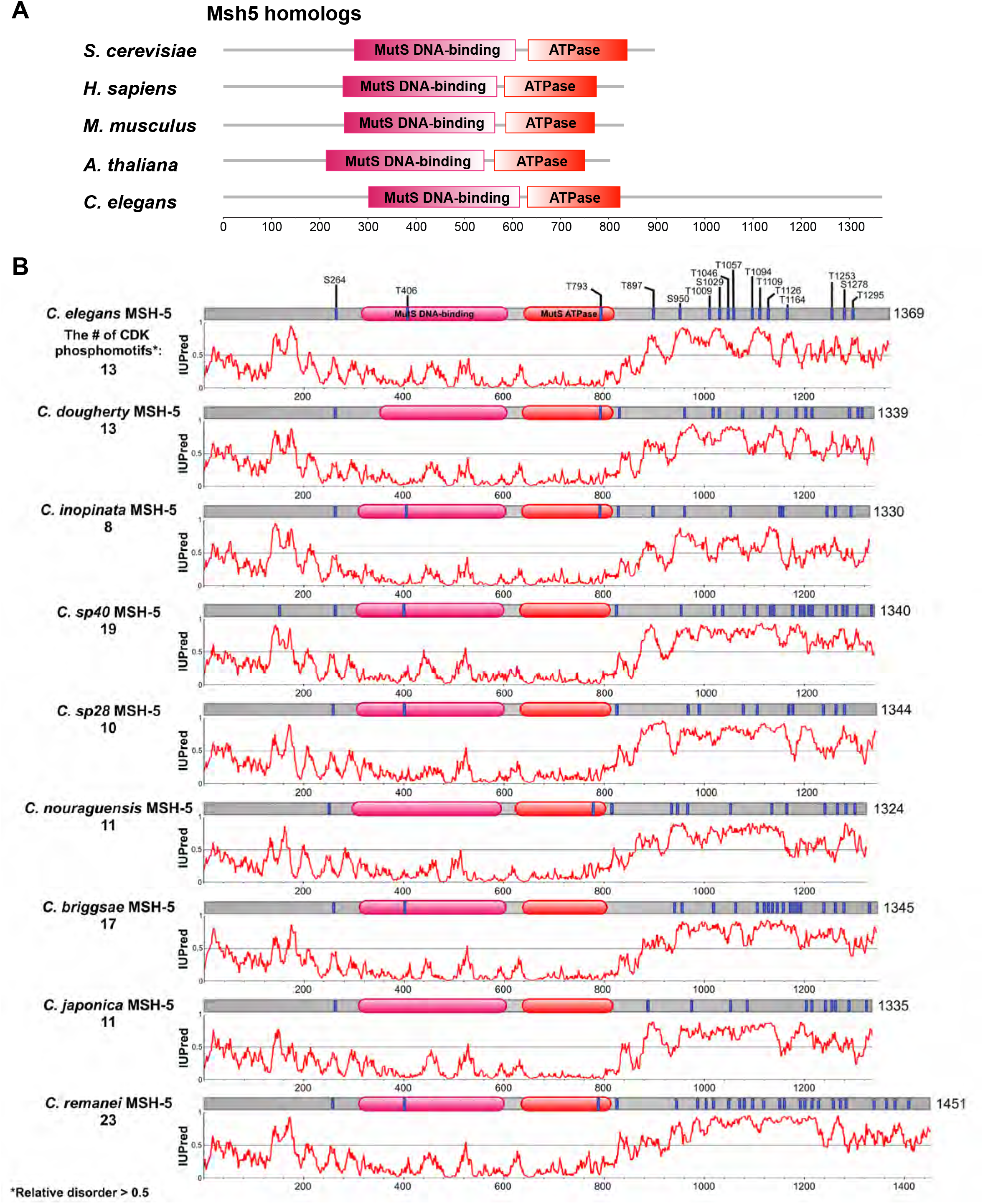
The domain structure of Msh5 orthologs and the distribution of CDK consensus sites in related *Caenorhabditis* species. (A) Schematic showing the domain structure of Msh5 orthologs from indicated eukaryotic species. (B) Schematic showing predicted CDK phosphorylation sites within MSH-5 orthologs from related *Caenorhabditis* species. The IUPred disorder plots are shown below. The number of CDK consensus motifs within the disordered C-terminal tail is indicated on the left.

**Figure S5 (related to Figure 5).**
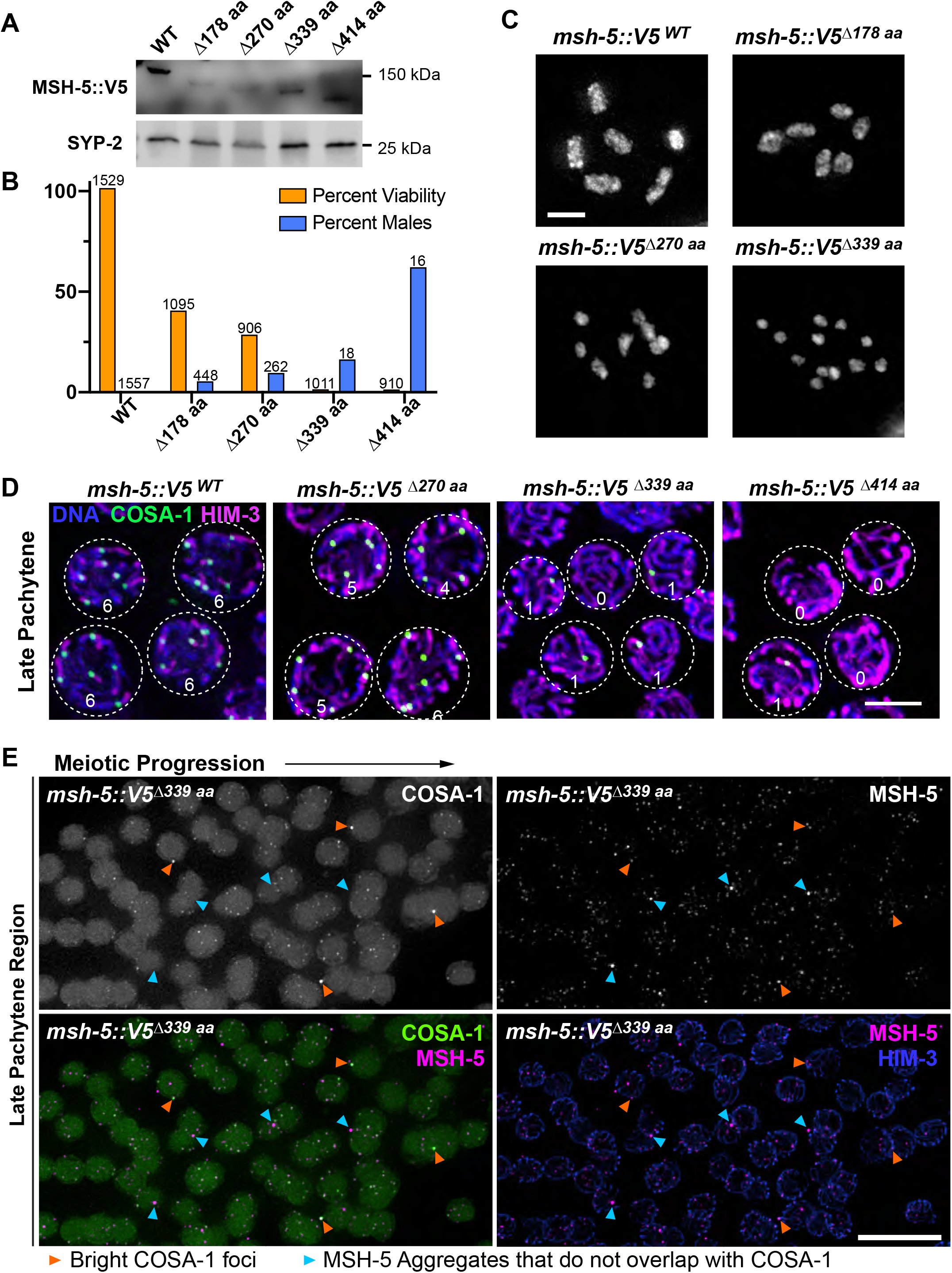
Analysis of *msh-5* C-terminal truncation mutants. (A) Immunoblot showing the expression of the MSH-5 C-terminal truncation proteins in whole worm lysates. SYP-2 was used as a loading control. (B) Graph showing the quantification of the percent viable self-progeny and males from indicated truncation mutants. Numbers of nuclei scored are shown on the top. (C) Oocyte nuclei at diakinesis from indicated genotypes were stained with DAPI. Scale bar, 3 μm. (D) Immunofluorescence images of late pachytene nuclei from the indicated genotypes showing DNA (blue), COSA-1 (green), and HIM-3 (magenta) staining. The boundary of individual nuclei is indicated by white dotted lines, and the number of COSA-1 foci in each nucleus is also shown. Scale bar, 3 μm. (E) Immunofluorescence of late pachytene region of a spread gonad from *msh-5::V5^Δ339aa^* showing the staining for COSA-1, MSH-5, and HIM-3. Scale bar, 10 μm.

**Figure S6 (related to Figure 6).**
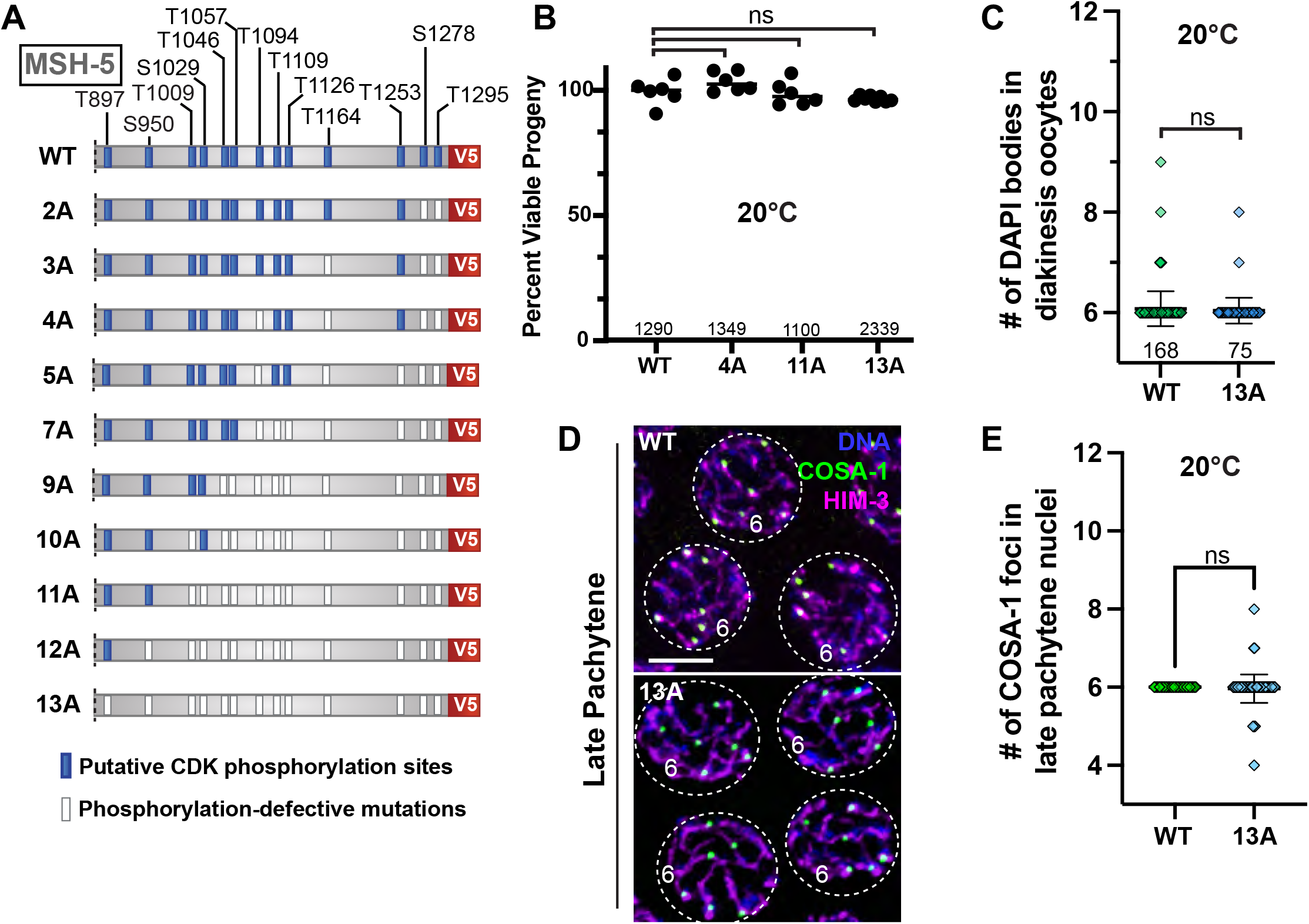
Mutation of all CDK consensus sites within MSH-5 C-terminal tail does not cause meiotic defects in an otherwise wild-type background. (A) Diagram of phosphorylation-defective *msh-5* alleles generated for this study. (B) Graph showing the viability of self-progeny from the indicated genotypes at 20°C. Numbers of eggs scored are indicated in the bottom. All *msh-5* mutants do not show significant differences from the wild-type. ns, not significant by ordinary one-way ANOVA (*msh-5::V5^WT^ vs. msh-5::V5^4A^,* p = 0.2526; *msh-5::V5^WT^ vs. msh-5::V5^11A^,* p = 0.9935, *msh-5::V5^WT^ vs. msh-5::V5^13A^,* p = 0.4988). (C) Graph showing quantification of DAPI bodies in diakinesis oocytes from *msh-5::V5^WT^* and *msh-5::V5^13A^* animals grown at 20°C. Mean ± SD is shown. Numbers of nuclei scored are indicated in the bottom. ns, not significant (p = 0.3240) by Mann-Whitney test. (D) Immunofluorescence images of late pachytene nuclei from *msh-5::V5^WT^* and *msh-5::V5^13A^* animals showing DNA (blue), COSA-1 (green), and HIM-3 (magenta) staining. The boundaries of individual nuclei are indicated by white dotted lines, and the number of COSA-1 foci per nucleus is indicated. Scale bar, 5 μm. (E) Graph showing the number of COSA-1 foci in late pachytene nuclei from *msh-5::V5^WT^* and *msh-5::V5^13A^* animals. Mean ± SD is shown. Numbers of nuclei scored are indicated in the bottom. ns, not significant (p = 0.1888) by Mann-Whitney test.

**Figure S7 (related to Figure 6).**
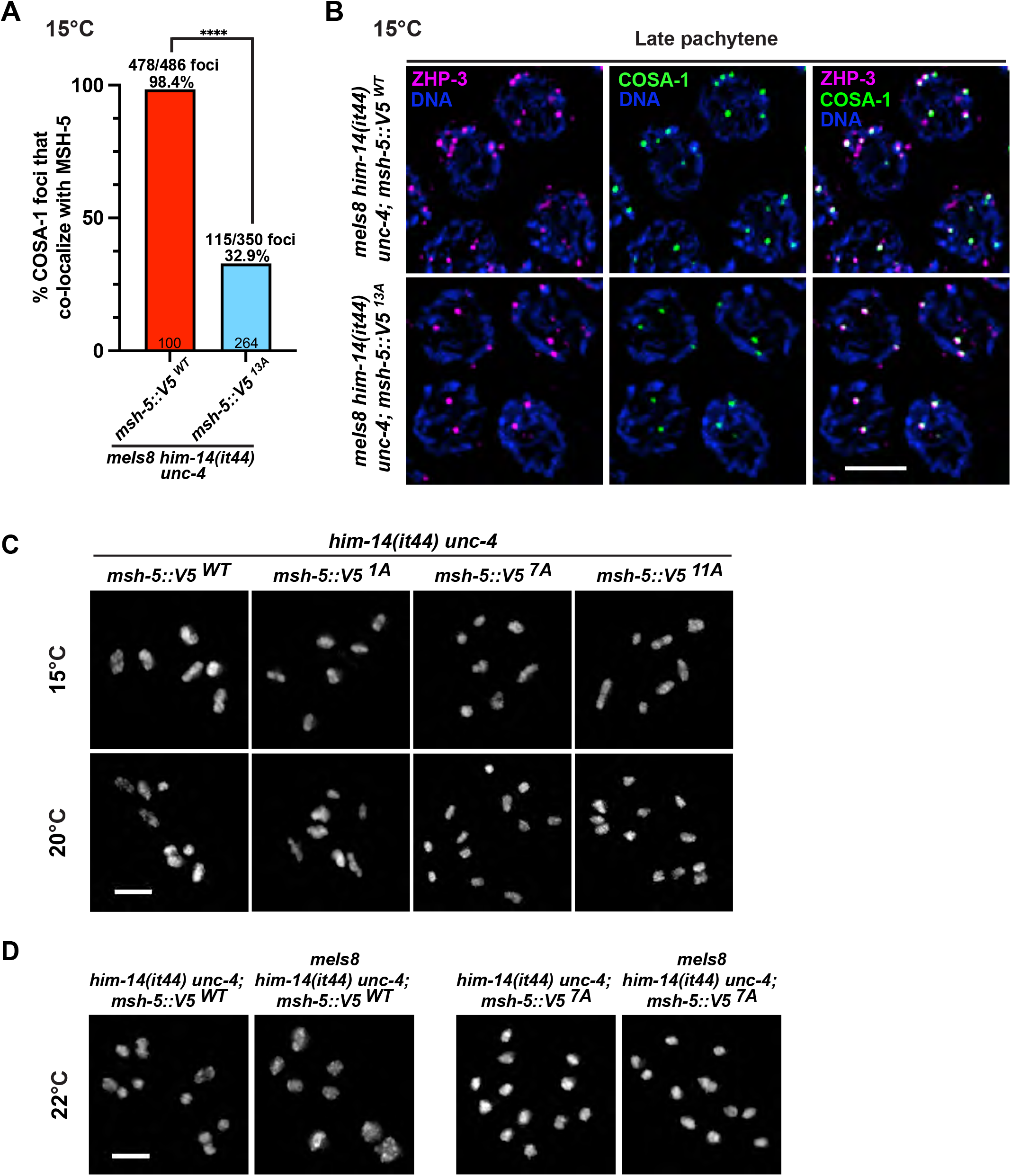
Localization of other pro-crossover factors in *him- 14(it44)* mutants harboring phospho-null mutations of *msh-5*. (A) Graph showing the percentage of COSA-1 foci co-localizing with MSH-5 from germline spreads of indicated genotypes grown at 15°C. Numbers of nuclei scored are shown below, and total foci counted are shown above. ****, p<0.0001 by two-tailed, unpaired t-test. (B) Immunofluorescence images of late pachytene nuclei from indicated genotypes grown at 15°C showing staining for DAPI (blue), ZHP-3 (magenta), and GFP::COSA-1 (green). Scale bar, 3 μm. (C) Oocyte nuclei at diakinesis from indicated genotypes grown at 15°C and 20°C were stained with DAPI. Scale bar, 3 μm. (D) DAPI-stained bodies in diakinesis oocytes from indicated genotypes at 22°C. Scale bar, 3 μm.

## Supplementary Tables

**Table S1.**
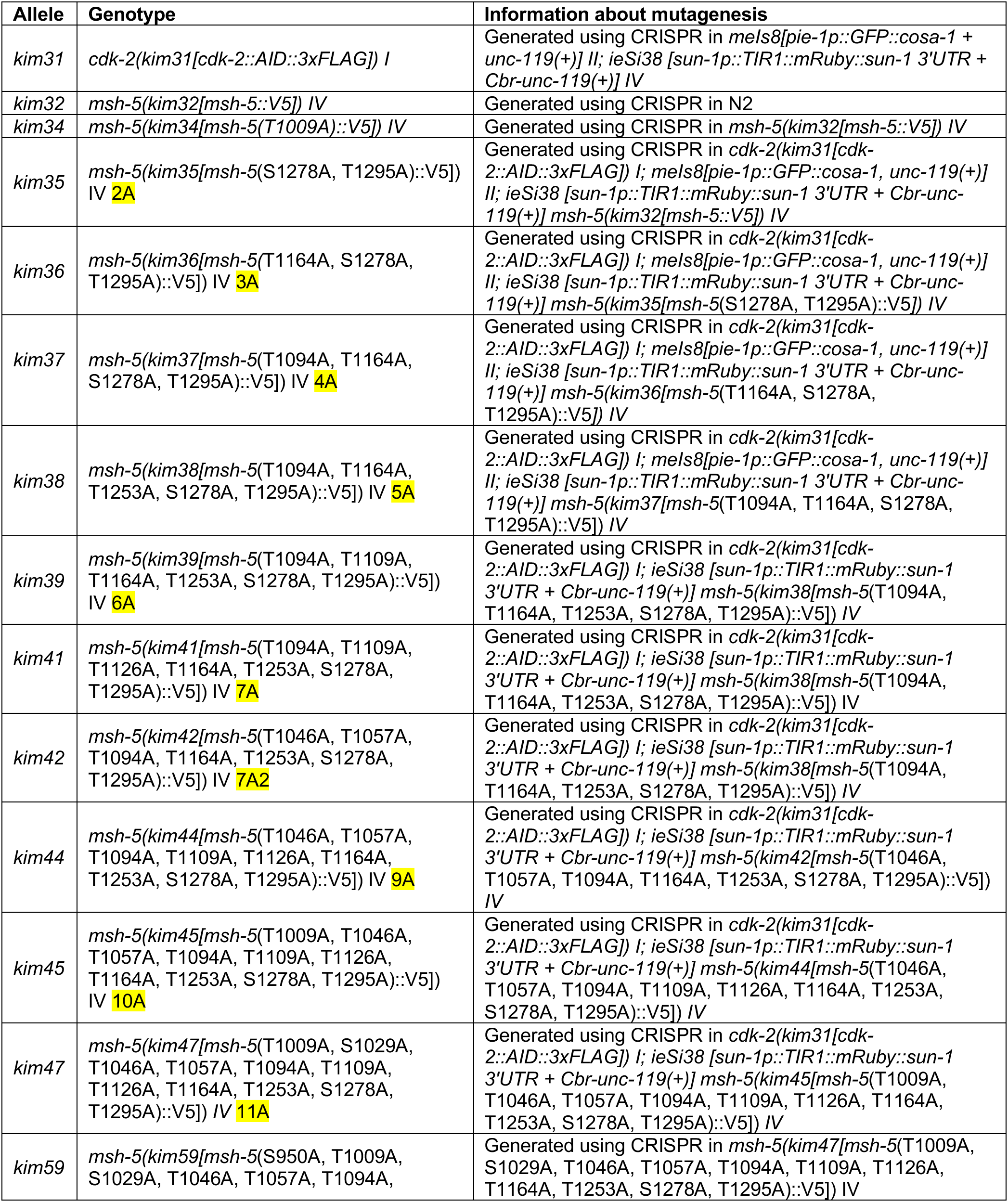
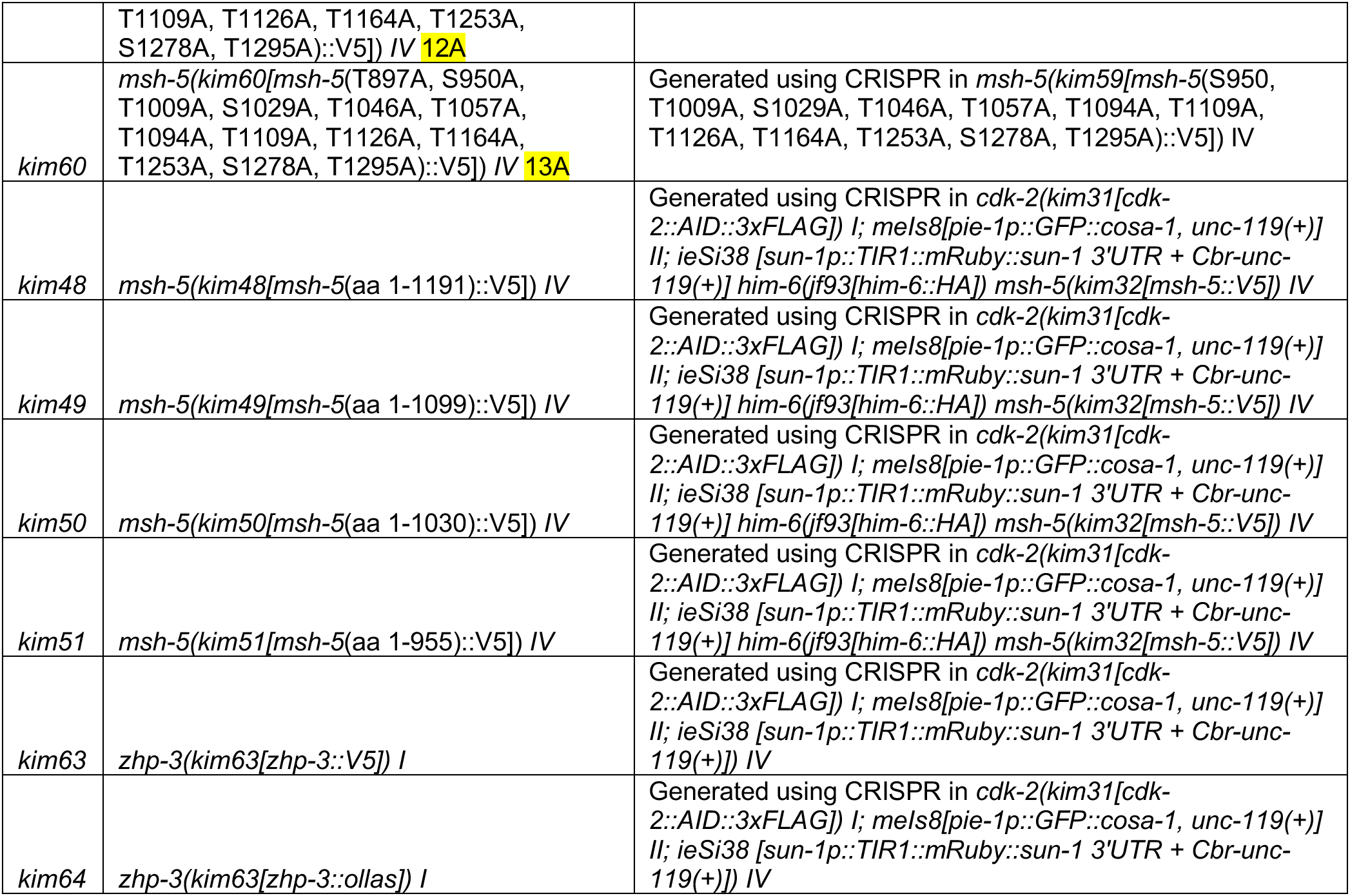
Alleles generated in this study.

**Table S2.**
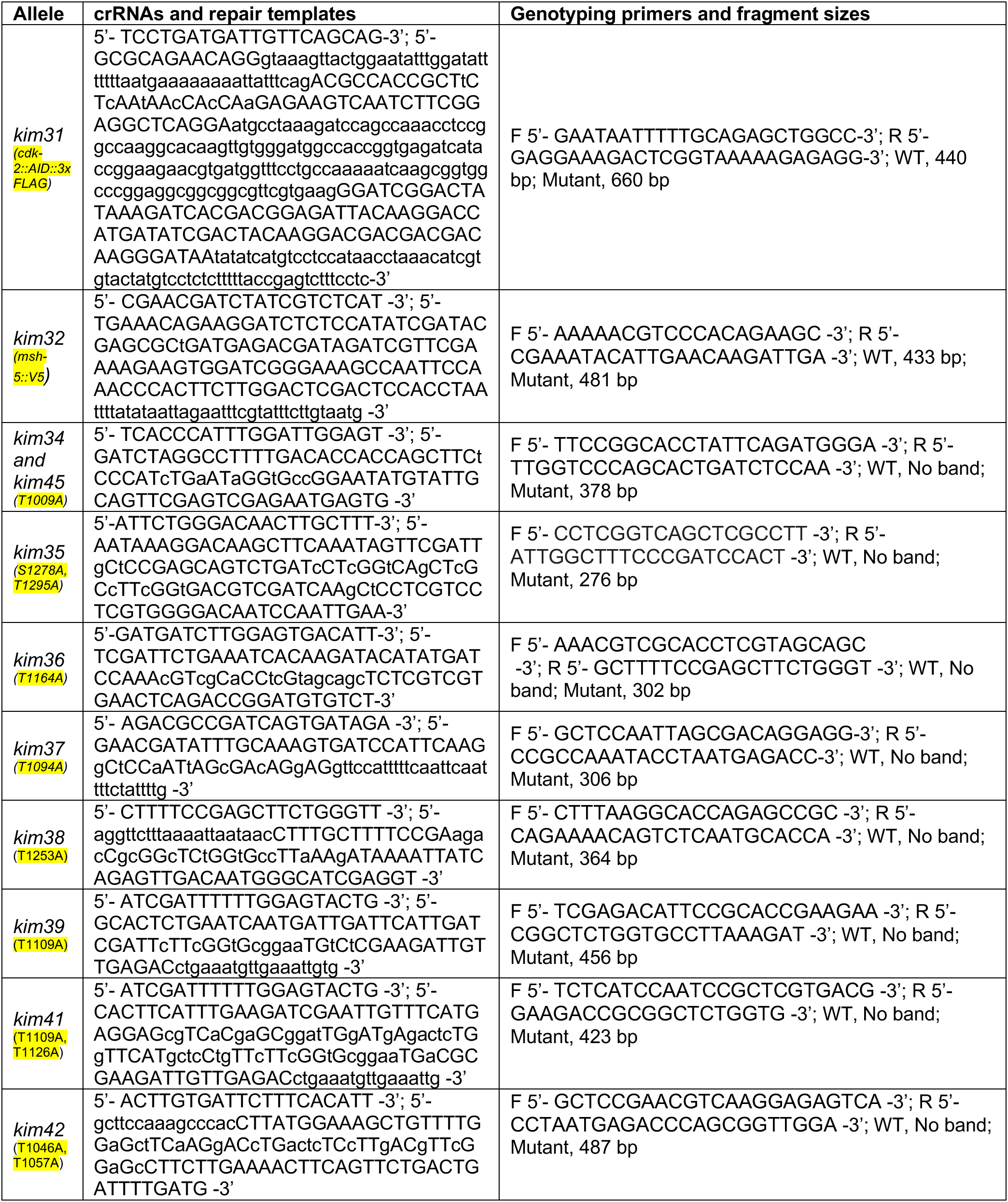
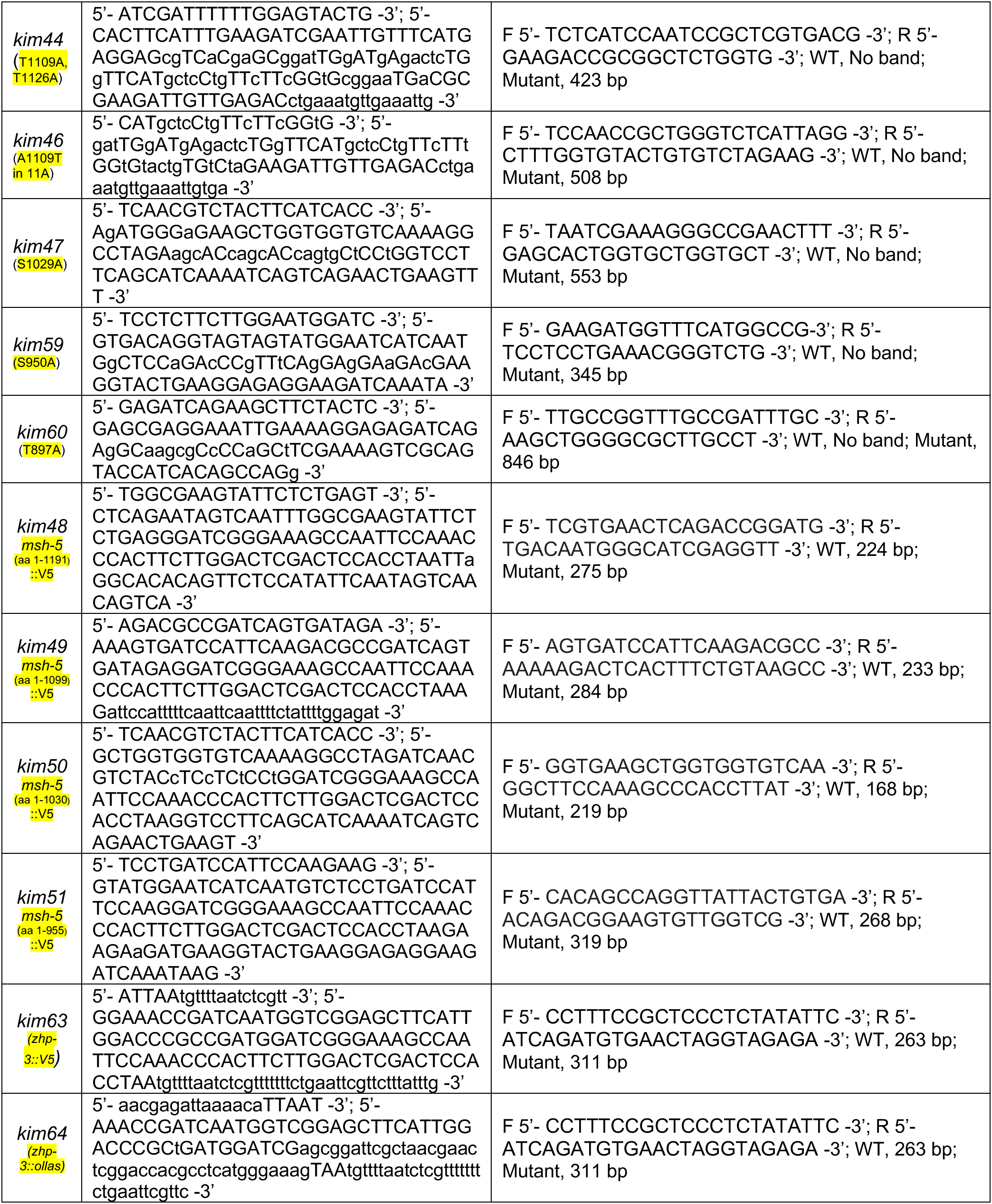
crRNAs, repair templates and genotyping primers for mutant alleles generated in this study.

**Table S3.** Strains used in this study.

| **Strains** | **Source** | **Strain #** |
| --- | --- | --- |
| N2 | Caenorhabditis Genetics Center | CB |
| *mels8[pie-1p::GFP::cosa-1, unc-119(+)] II* | Yokoo et al., 2012 | AV630 |
| *cdk-2(kim31[cdk-2::AID::3xFLAG]) I; mels8[pie-1p::GFP::cosa-1, unc-119(+)] II; ieSi38 [sun-1p::TIR1::mRuby::sun-1 3'UTR + Cbr-unc-119(+)] IV* | This study | YKIM110 |
| *him-6(jf93[him-6::HA]) IV* | Jagut et al., 2016 | UV116 |
| *cdk-2(kim31[cdk-2::AID::3xFLAG]) I; mels8[pie-1p::GFP::cosa-1, unc-119(+)] II; ieSi38 [sun-1p::TIR1::mRuby::sun-1 3'UTR + Cbr-unc-119(+)] him-6(jf93[him-6::HA]) msh-5(kim32[msh-5::V5]) IV* | This study | YKM762 |
| *cdk-2(kim31[cdk-2::AID::3xFLAG]) I; mels8[pie-1p::GFP::cosa-1, unc-119(+)] II; ieSi38 [sun-1p::TIR1::mRuby::sun-1 3'UTR + Cbr-unc-119(+)] msh-5(kim32[msh-5::V5]) IV* | This study | YKM306 |
| *msh-5(kim32[msh-5::V5]) IV* | This study | YKM154 |
| *him-14(it44) unc-4(e120)/mnC1 II* | Girard et al., 2021 | KK323 |
| *mels8[pie-1p::GFP::cosa-1, unc-119(+)] him-14(it44) unc-4(e120)/mnC1 II; oxTi602 IV* | This study | AV1176 |
| *him-14(it44) unc-4(e120)/mnC1 II; msh-5(kim32[msh-5::V5]) IV* | This study | AV1172 |
| *mels8[pie-1p::GFP::cosa-1, unc-119(+)] him-14(it44) unc-4(e120)/mnC1 II; msh-5(kim32[msh-5::V5]) IV* | This study | AV1171 |
| *msh-5(kim34[msh-5(T1009A)::V5]) IV* | This study | YKM233 |
| *him-14(it44) unc-4(e120)/mnC1 II; msh-5(kim34[msh-5(T1009A)::V5]) IV* | This study | AV1166 |
| *mels8[pie-1p::GFP::cosa-1, unc-119(+)] him-14(it44) unc-4(e120)/mnC1 II; msh-5(kim34[msh-5(T1009A)::V5]) IV* | This study | AV1165 |
| *cdk-2(kim31[cdk-2::AID::3xFLAG]) I; mels8[pie-1p::GFP::cosa-1, unc-119(+)] II; ieSi38 [sun-1p::TIR1::mRuby::sun-1 3'UTR + Cbr-unc-119(+)] msh-5(kim35[msh-5(S1278A, T1295A)::V5]) IV* | This study | YKM398 |
| *cdk-2(kim31[cdk-2::AID::3xFLAG]) I; mels8[pie-1p::GFP::cosa-1, unc-119(+)] II; ieSi38 [sun-1p::TIR1::mRuby::sun-1 3'UTR + Cbr-unc-119(+)] msh-5(kim36[msh-5(T1164A, S1278A, T1295A)::V5]) IV* | This study | YKM459 |
| *cdk-2(kim31[cdk-2::AID::3xFLAG]) I; mels8[pie-1p::GFP::cosa-1, unc-119(+)] II; ieSi38 [sun-1p::TIR1::mRuby::sun-1 3'UTR + Cbr-unc-119(+)] msh-5(kim37[msh-5(T1094A, T1164A, S1278A, T1295A)::V5]) IV* | This study | YKM477 |
| *cdk-2(kim31[cdk-2::AID::3xFLAG]) I; mels8[pie-1p::GFP::cosa-1, unc-119(+)] II; ieSi38 [sun-1p::TIR1::mRuby::sun-1 3'UTR + Cbr-unc-119(+)] msh-5(kim38[msh-5(T1094A, T1164A, T1253A, S1278A, T1295A)::V5]) IV* | This study | YKM460 |
| *cdk-2(kim31[cdk-2::AID::3xFLAG]) I; ieSi38 [sun-1p::TIR1::mRuby::sun-1 3'UTR + Cbr-unc-119(+)] msh-5(kim38[msh-5(T1094A, T1164A, T1253A, S1278A, T1295A)::V5]) IV* | This study | YKM478 |
| *cdk-2(kim31[cdk-2::AID::3xFLAG]) I; ieSi38 [sun-1p::TIR1::mRuby::sun-1 3'UTR + Cbr-unc-119(+)] msh-5(kim39[msh-5(T1094A, T1109A, T1164A, T1253A, S1278A, T1295A)::V5]) IV* | This study | YKM535 |
| *cdk-2(kim31[cdk-2::AID::3xFLAG]) I; ieSi38 [sun-1p::TIR1::mRuby::sun-1 3'UTR + Cbr-unc-119(+)] msh-5(kim41[msh-5(T1094A, T1109A, T1126A, T1164A, T1253A, S1278A, T1295A)::V5]) IV* | This study | YKM532 |
| *him-14(it44) unc-4(e120)/mnC1 II; msh-5(kim41[msh-5(T1094A, T1109A, T1126A, T1164A, T1253A, S1278A, T1295A)::V5]) IV* | This study | AV1168 |
| *mels8[pie-1p::GFP::cosa-1, unc-119(+)] him-14(it44) unc-4(e120)/mnC1 II; msh-5(kim41[msh-5(T1094A, T1109A, T1126A, T1164A, T1253A, S1278A, T1295A)::V5]) IV* | This study | AV1167 |
| *cdk-2(kim31[cdk-2::AID::3xFLAG]) I; ieSi38 [sun-1p::TIR1::mRuby::sun-1 3'UTR + Cbr-unc-119(+)] msh-5(kim42[msh-5(T1046A, T1057A, T1094A, T1164A, T1253A, S1278A, T1295A)::V5]) IV* | This study | YKM556 |
| *cdk-2(kim31[cdk-2::AID::3xFLAG]) I; ieSi38 [sun-1p::TIR1::mRuby::sun-1 3'UTR + Cbr-unc-119(+)] msh-5(kim44[msh-5(T1046A, T1057A, T1094A, T1109A, T1126A, T1164A, T1253A, S1278A, T1295A)::V5]) IV* | This study | YKM505 |
| *cdk-2(kim31[cdk-2::AID::3xFLAG]) I; ieSi38 [sun-1p::TIR1::mRuby::sun-1 3'UTR + Cbr-unc-119(+)] msh-5(kim45[msh-5(T1009A, T1046A, T1057A, T1094A, T1109A, T1126A, T1164A, T1253A, S1278A, T1295A)::V5]) IV* | This study | YKM506 |
| *cdk-2(kim31[cdk-2::AID::3xFLAG]) I; ieSi38 [sun-1p::TIR1::mRuby::sun-1 3'UTR + Cbr-unc-119(+)] msh-5(kim47[msh-5(T1009A, S1029A, T1046A, T1057A, T1094A, T1109A, T1126A, T1164A, T1253A, S1278A, T1295A)::V5]) IV* | This study | YKM507 |
| *msh-5(kim47[msh-5(T1009A, S1029A, T1046A, T1057A, T1094A, T1109A, T1126A, T1164A, T1253A, S1278A, T1295A)::V5]) IV* | This study | YKM655 |
| *him-14(it44) unc-4(e120)/mnC1 II; msh-5(kim47[msh-5(T1009A, S1029A, T1046A, T1057A, T1094A, T1109A, T1126A, T1164A, T1253A, S1278A, T1295A)::V5]) IV* | This study | AV1174 |
| *mels8[pie-1p::GFP::cosa-1, unc-119(+)] him-14(it44) unc-4(e120)/mnC1 II; msh-5(kim47[msh-5(T1009A, S1029A, T1046A, T1057A, T1094A, T1109A, T1126A, T1164A, T1253A, S1278A, T1295A)::V5]) IV* | This study | AV1173 |
| *msh-5(kim59[msh-5(S950A, T1009A, S1029A, T1046A, T1057A, T1094A, T1109A, T1126A, T1164A, T1253A, S1278A, T1295A)::V5]) IV* | This study | YKM685 |
| *msh-5(kim60[msh-5(T897A, S950A, T1009A, S1029A, T1046A, T1057A, T1094A, T1109A, T1126A, T1164A, T1253A, S1278A, T1295A)::V5]) IV* | This study | YKM692 |
| *rmels8[pie-1p::GFP::cosa-1, unc-119(+)] II; msh-5(kim60[msh-5(T897A, S950A, T1009A, S1029A, T1046A, T1057A, T1094A, T1109A, T1126A, T1164A, T1253A, S1278A, T1295A)::V5]) IV* | This study | YKM719 |
| *him-14(it44) unc-4(e120)/mnC1 II; msh-5(kim60[msh-5(T897A, S950A, T1009A, S1029A, T1046A, T1057A, T1094A, T1109A, T1126A, T1164A, T1253A, S1278A, T1295A)::V5]) IV* | This study | AV1170 |
| *mels8[pie-1p::GFP::cosa-1, unc-119(+)] him-14(it44) unc-4(e120)/mnC1 II; msh-5(kim60[msh-5(T897A, S950A, T1009A, S1029A, T1046A, T1057A, T1094A, T1109A, T1126A, T1164A, T1253A, S1278A, T1295A)::V5]) IV* | This study | AV1169 |
| *cdk-2(kim31[cdk-2::AID::3xFLAG]) I; mels8[pie-1p::GFP::cosa-1, unc-119(+)] II; ieSi38 [sun-1p::TIR1::mRuby::sun-1 3'UTR + Cbr-unc-119(+)] him-6(jf93[him-6::HA]) msh-5(kim48[msh-5(aa 1-1191)::V5]) IV* | This study | YKM357 |
| *cdk-2(kim31[cdk-2::AID::3xFLAG]) I; mels8[pie-1p::GFP::cosa-1, unc-119(+)] II; ieSi38 [sun-1p::TIR1::mRuby::sun-1 3'UTR + Cbr-unc-119(+)] him-6(jf93[him-6::HA]) msh-5(kim49[msh-5(aa 1-1099)::V5]) IV* | This study | YKM454 |
| *cdk-2(kim31[cdk-2::AID::3xFLAG]) I; mels8[pie-1p::GFP::cosa-1, unc-119(+)] II; ieSi38 [sun-1p::TIR1::mRuby::sun-1 3'UTR + Cbr-unc-119(+)] him-6(jf93[him-6::HA]) msh-5(kim50[msh-5(aa 1-1030)::V5]) IV/nT1 [unc-?(n754) let-?] (IV;V)* | This study | YKM401 |
| *cdk-2(kim31[cdk-2::AID::3xFLAG]) I; mels8[pie-1p::GFP::cosa-1, unc-119(+)] II; ieSi38 [sun-1p::TIR1::mRuby::sun-1 3'UTR + Cbr-unc-119(+)] him-6(jf93[him-6::HA]) msh-5(kim51[msh-5(aa 1-955)::V5]) IV/nT1 [unc-?(n754) let-?] (IV;V)* | This study | YKM402 |
| *cdk-2(kim31[cdk-2::AID::3xFLAG]) zhp-3(kim63[zhp-3::V5]) I; mels8[pie-1p::GFP::cosa-1, unc-119(+)] II; ieSi38 [sun-1p::TIR1::mRuby::sun-1 3'UTR + Cbr-unc-119(+)] IV* | This study | YKM165 |
| *cdk-2(kim31[cdk-2::AID::3xFLAG]) zhp-3(kim63[zhp-3::ollas]) I; mels8[pie-1p::GFP::cosa-1, unc-119(+)] II; ieSi38 [sun-1p::TIR1::mRuby::sun-1 3'UTR + Cbr-unc-119(+)] msh-5(kim32[msh-5::V5]) IV* | This study | YKM760 |
| *cdk-2(kim31[cdk-2::AID::3xFLAG]) zhp-3(kim63[zhp-3::ollas]) I; mels8[pie-1p::GFP::cosa-1, unc-119(+)] II; ieSi38 [sun-1p::TIR1::mRuby::sun-1 3'UTR + Cbr-unc-119(+)] him-6(jf93[him-6::HA]) msh-5(kim50[msh-5(aa 1-1030)::V5]) IV/nT1 [unc-?(n754) let-?] (IV;V)* | This study | YKM761 |

**Table S4.**
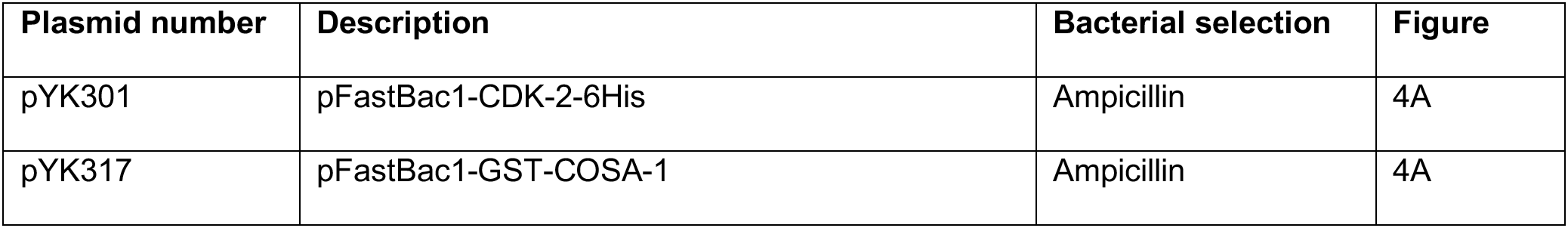
Plasmids used in this study.

Plasmids used in this study
| Plasmid number | Description | Bacterial selection | Figure |
| --- | --- | --- | --- |
| pYK301 | pFastBac1-CDK-2-6His | Ampicillin | 4A |
| pYK317 | pFastBac1-GST-COSA-1 | Ampicillin | 4A |

